# MEK1/2 inhibition transiently alters the tumor immune microenvironment to enhance immunotherapy efficacy against head and neck cancer

**DOI:** 10.1101/2021.08.22.457244

**Authors:** Manu Prasad, Jonathan Zorea, Sankar Jagadeeshan, Avital Shnerb, Jebrane Bouaoud, Lucas Michon, Ofra Novoplansky, Mai Badarni, Limor Cohen, Ksenia Yagodayev, Sapir Tzadok, Barak Rotblat, Libor Brezina, Andreas Mock, Andy Karabajakian, Jérôme Fayette, Idan Cohen, Tomer Cooks, Irit Allon, Orr Dimitstei, Benzion Joshua, Dexin Kong, Elena Voronov, Maurizio Scaltriti, Yaron Carmi, Jochen Hess, Luc G.T. Morris, Pierre Saintigny, Moshe Elkabets

**Affiliations:** The Shraga Segal Department of Microbiology, Immunology, and Genetics, Ben-Gurion University of the Negev; Beer-Sheva, Israel; Faculty of Health Sciences, Ben-Gurion University of the Negev; Beer-Sheva, Israel; Université Claude Bernard Lyon, INSERM 1052, CNRS 5286, Centre Léon Bérard, Centre de Recherche en Cancérologie de Lyon, Lyon 69373, France.; Department of Translational Medicine Oncology, Centre Léon Bérard; Lyon 69373, France.; Department of Life Sciences, Ben-Gurion University of the Negev; Beer-Sheva, Israel; The National Institute for Biotechnology in the Negev, Ben-Gurion University of the Negev; Beer-Sheva, Israel; Department of Medical Oncology, National Center for Tumor Diseases (NCT) Heidelberg, Heidelberg University Hospital, Heidelberg, Germany; Division of Translational Medical Oncology, NCT Heidelberg, German Cancer Center (DKFZ), Heidelberg, Germany; Department of Medical Oncology, Centre Léon Bérard; Lyon 69373, France.; Institute of Pathology, Barzilai University Medical Center; Ashkelon, Israel.; Department of Otolaryngology-Head & Neck Surgery, Soroka University Medical Center; Beer-Sheva, Israel.; Department of Otorhinolaryngology and Head & Neck Surgery, Barzilai Medical Center; Ashkelon, Israel.; School of Pharmaceutical Sciences, Tianjin Medical University; Tianjin 300070, China; Human Oncology and Pathogenesis Program, Memorial Sloan Kettering Cancer Center; New York, USA.; Department of Pathology, Sackler School of Medicine, Tel Aviv University, Tel Aviv, Israel.; Section Experimental and Translational Head and Neck Oncology, Department of Otolaryngology, Head and Neck Surgery, University Hospital Heidelberg, Heidelberg, Germany; Research Group Molecular Mechanisms of Head and Neck Tumors, German Cancer Research Center (DKFZ); Heidelberg, Germany.; Department of Surgery, Memorial Sloan Kettering Cancer Center; New York, USA.

**Keywords:** Head and neck cancer, tumor-microenvironment, tumor-immunity, chemokines, immunotherapy, targeted therapy, MEK1/2, anti-PD-1

## Abstract

Although the mitogen-activated protein kinases (MAPK) pathway is hyperactive in head and neck cancer (HNC), inhibition of MEK1/2 in HNC patients has not shown clinically meaningful activity. Using pre-clinical HNC models, we demonstrated that treatment with the MEK1/2 blocker trametinib delays HNC initiation and progression by reducing tumor cell proliferation and enhancing the anti-tumor immunity of CD8^+^ T cells. Further activation of CD8^+^ T cells by supplementation with anti-programmed death-1 (αPD-1) antibody eliminated tumors and induced an immune memory in the cured mice. Mechanistically, an early response to trametinib treatment sensitized tumors to αPD-1-supplementation by attenuating the expression of tumor-derived colony-stimulating factor-1 (CSF-1), which reduced the abundance of two CSF-1R^+^CD11c^+^ myeloid-derived suppressor cell (MDSC) populations in the tumor microenvironment (TME). In contrast, prolonged treatment with trametinib abolished the anti-tumor activity of αPD-1, because tumor cells undergoing the epithelial to mesenchymal transition (EMT) in response to trametinib restored CSF-1 expression and re-created an immune-suppressive TME. These findings provide the rationale for testing the trametinib/αPD-1 combination in HNC and highlight the importance of sensitizing tumors to immunotherapies by using targeted therapies to interfere with the host-tumor interaction.

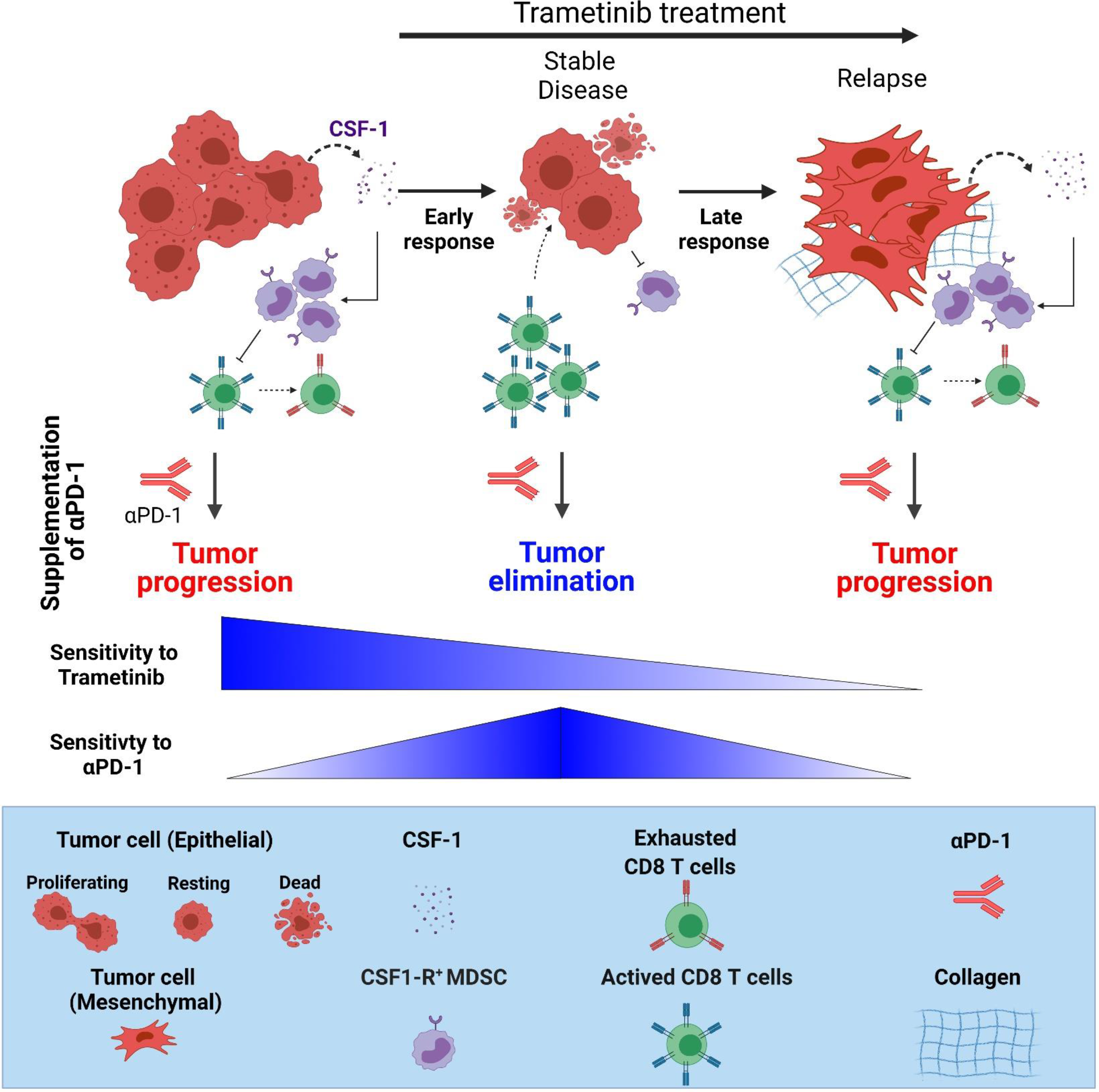
Graphical abstract.

## Introduction

Genomic alterations in genes of the mitogen-activated protein kinase (MAPK) pathway, or activation of receptor tyrosine kinases (RTK), induce constant activation of MEK-ERK signaling, which regulates cell proliferation, survival, migration, and transformation (as reviewed in (1, 2)). Since hyperactivation of the MAPK pathway is involved in the pathogenesis of head and neck cancer (HNC) ((3, 4) and reviewed in (5)), blockade of the MAPK pathway has been studied as a therapeutic approach for counteracting HNC progression. For example, cetuximab, an anti-epidermal growth factor receptor (EGFR) antibody that blocks the MAPK pathway, showed a significant but transient anti-tumor effect in HNC patients (6). Currently, multiple clinical trials are testing MAPK/RTK inhibitors in combination with different modalities in patients with HNC. Trametinib, a MEK1/2 kinase inhibitor approved for treatment of MAPK-driven melanoma and non-small cell lung cancer, exhibited a favorable but transient anti-tumor effect in HNC patients with oral squamous cell carcinoma (7). Trametinib, and other blockers of the MAPK pathway, act by arresting the growth of MAPK-driven tumor cells, but they also affect the host immunity and the heterogeneity of the tumor microenvironment (TME) (8–10). The TME is composed of different stromal and immune cell types, which interact with each other and with the malignant cells and thus determine tumor progression (reviewed in (11, 12)). The complex communication network between malignant cells and the TME is mediated, in part, by the tumor cell-derived cytokines and chemokines that regulate the heterogeneity of the TME and support tumor progression and/or resistance to therapy ((13) and reviewed in (14–16)). Using pre-clinical HNC models, it was shown that treatment of tumor-bearing mice with trametinib induced CD8^+^ T cell infiltration into the TME and upregulated expression of programmed death-ligand 1 (PD-L1) by the tumor cells (17). It was also shown that combined treatment with trametinib and anti-PD-L1 antibody (αPD-L1) arrested tumor growth in the mice (17). While the above lines of evidence support the involvement of MAPK inhibitors in regulating the heterogeneity of TME and thus in determining anti-tumor immunity, the mechanisms of the communication network between the malignant cells and the TME are yet to be fully understood.

Here we found that trametinib treatment delays tumor initiation and progression of MAPK-pathway mutated HNC, while altering the heterogeneity of TME, in part via downregulating the expression of tumor-derived colony-stimulating factor-1 (CSF-1). Tumor-derived CSF-1 controlled the quantity of CSF-1R^+^CD11c^+^ MDSCs and the infiltration of activated CD8^+^ T cells into the tumor, which subsequently affected the sensitivity to the FDA-approved immunotherapy for HNC, anti-PD-1 (αPD-1) (18). We showed that the timing of αPD-1 supplementation to trametinib treatment is crucial, as tumor elimination occurred only when αPD-1 supplementation was administered in the time window during which trametinib treatment had temporarily reduced CSF-1 expression and induced an immune active TME. This transient immune activation, reflected by low CSF1 expression and high CD8A expression, was positively associated with a clinical benefit for HNC patients treated with αPD1.

## Results

### MAPK pathway blockade with trametinib delays HNC initiation and progression

We first established the frequency of genomic alterations in the genes associated with the MAPK pathway in multiple HNC cohorts by interrogating available databases and found that between 5% -50% of genes are altered (Supplementary Fig. S1A, and Supplementary Table S1). Targeted sequencing of 1680 HNC patients (GENIE cohort), and whole-exome sequencing of 676 HNC patients showed that the frequency of mutations in MAPK related genes was 13% and 29%, respectively (Supplementary Fig. S1B, C). Moreover, inference of pathway activity based on RNA sequencing (RNA-seq) data from The Cancer Genome Atlas (TCGA)-HNC revealed significant associations of MAPK pathway hyperactive HNC with a worse prognosis, a larger tumor size, a basal classified signature, and human papillomavirus 16-negative (HPV16^-^) status as compared to patients with low MAPK pathway activity (Supplementary Fig. S1D, and Supplementary Tables S2 and S3). In addition, the expression of ERK2, a key component of the MAPK signaling pathway that is encoded by the *MAPK1* gene, is upregulated in HNC tumors compared to normal tissue, and there is an association between the extent of upregulation and the tumor grade (Supplementary Fig. S1E, F) (UALCAN(19) -TCGA data). Lastly, analysis of the Cancer Cell Line Encyclopedia (CCLE) and the Sanger cell line datasets showed that HNC cell lines are sensitive to trametinib (Supplementary Fig. S1G), a conclusion supported by the recent publication of Lepikhova et al.(20).

Following the above described analysis of the published data, we assessed the efficacy of trametinib in two independent HNC patient-derived xenografts (PDXs) in which trametinib treatment either stabilized or significantly delayed tumor growth (Fig. 1A). Immunohistochemistry (IHC) analysis of the PDXs showed that trametinib reduced cell proliferation (Ki67 staining) and phosphorylated ERK1 and 2 (pERK1/2) levels as a ’readout’ of MAPK pathway activation (Fig. 1B). To investigate the impact of MAPK pathway activation on HNC initiation and progression in pre-clinical models, we initially determined pERK1/2 levels at various stages of oral carcinogenesis induced by the mutagenic compound 4-nitroquinoline-1-oxide (4NQO)(21) in C57BL/6J wild-type (WT) mice. Histopathological analysis of the tongues of 4NQO-treated mice showed a significant increase in pERK1/2 levels from lower pathological grades (11) (dysplasia) to higher pathological stages, namely, advanced squamous cell carcinoma (SCC) (Fig. 1C). Higher pERK1/2 levels and more advanced pathological grades were associated with increased cell proliferation, as indicated by Ki67 staining (Supplementary Fig. S1H). To explore whether targeting the MAPK pathway could be used for chemoprevention, we exposed WT mice to 4NQO for 12 weeks and then treated them with either trametinib or vehicle (Fig. 1D). The vehicle-treated mice started to develop physical signs of tumor progression, manifested by weight loss and swallowing difficulties after 14 weeks (∼100 days) and began to die after 160 days (Fig. 1D). In contrast, trametinib-treated mice started to show signs of disease onset after as long as 250 days. All vehicle-treated mice died within 220 days, while trametinib-treated mice survived significantly longer, with an extension of 100 days in median survival (Fig. 1D). An independent experiment in which we monitored disease progression and histopathology of mouse tongues on day 150 further confirmed that trametinib delayed tumor development (Supplementary Fig. S1I). In the vehicle-treated group, we detected 4–6 macroscopic lesions on the tongue of each mouse, whereas in the trametinib-treated group only 1–2 lesions per mouse were observed (Supplementary Fig. S1I). Moreover, mice treated with trametinib exhibited a less invasive phenotype, as determined by the depth of Invasion (DOI) and reduced hyperplasia of the basal layer of the tongue (22) (Supplementary Fig. S1J). Analysis of the tongue lesions showed the development of carcinoma in situ and advanced SCC in 55% of vehicle-treated mice, compared to only 5% of trametinib-treated mice (Fig. 1E).

**Figure 1.**
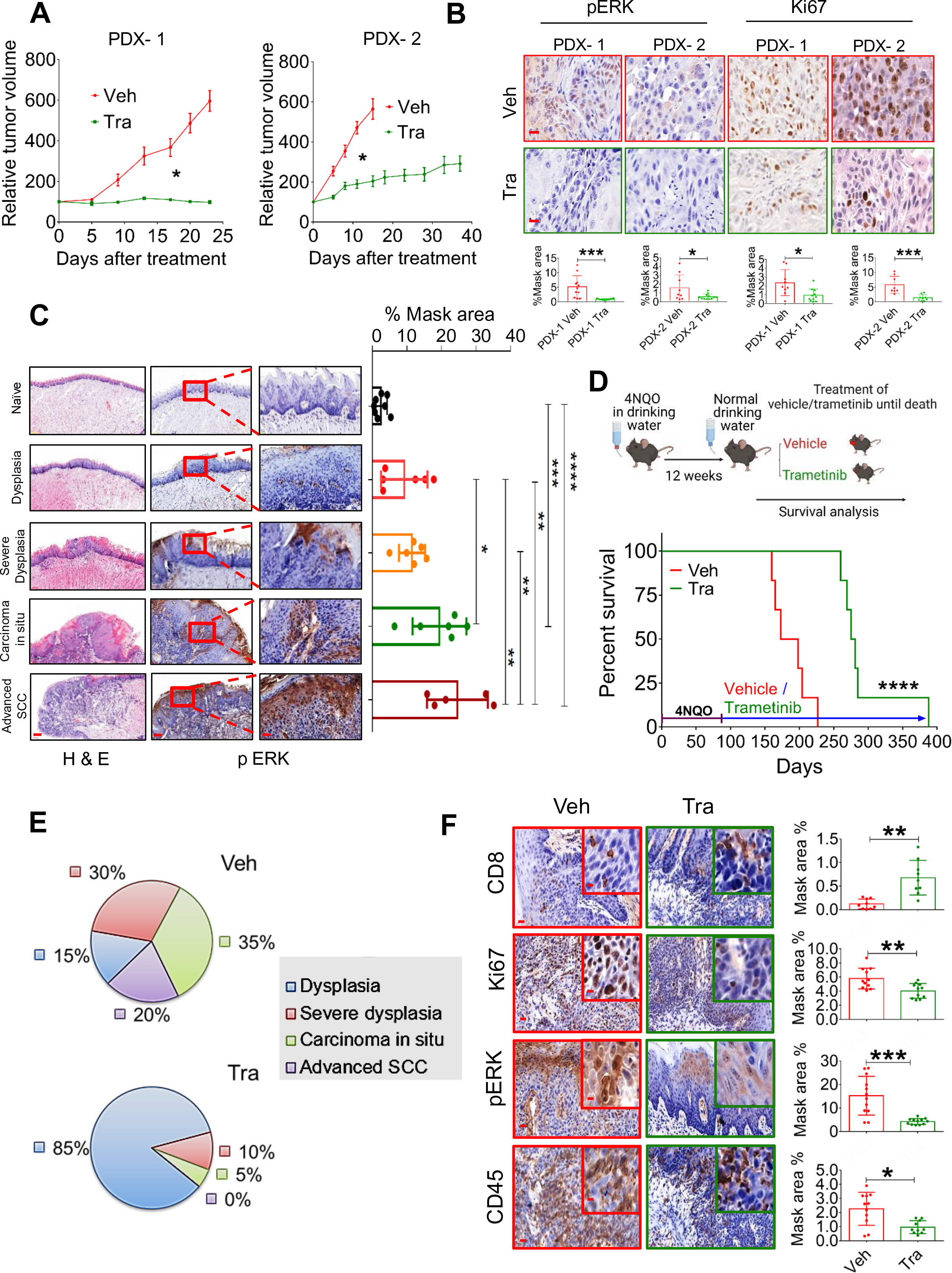
MAPK pathway is hyperactive in HNC, and trametinib induced MAPK pathway blockage delays HNC initiation and progression. (**A)** Relative tumor growth of two HNC-PDXs after treatment with trametinib. Values in the graphs are mean tumor volumes ± SEM. (**B)** Top –IHC staining of pERK and Ki67 in tissues of two HNC-PDXs after treatment with trametinib. Bottom – quantification of staining performed using 3DHISTECH software HistoQuant (n > 10) (scale bar: 10 µm). (**C)** H&E (left) and pERK (right) staining at various stages of oral (tongue) carcinogenesis induced by 4NQO in C57BL/6Jmice. % mask area is (**D)** Top – .shown on the right. Scale bars: 200 µm (left); 100 µm (middle); and 20 µm (right) Scheme of the experimental setting investigating the survival of 4NQO-treated mice subsequently treated with trametinib or vehicle. Bottom – Survival rates of immunocompetent C57BL/6J mice (n = 6) in a 4NQO cancer model following daily treatments with vehicle or trametinib (1 mg/kg/day). (**E)** Pie diagrams showing the percentages of different cancer grades in vehicle- and trametinib-treated mice. (**F)** IHC images showing the infiltration of CD45^+^ cells, CD8^+^ T cells, and the expression of pERK and Ki67 in the tongues of mice treated with vehicle or trametinib (scale bars: 20 µm; inset 10 µm). For statistics, an unpaired 2-sided *t*-test or one-way ANOVA was performed. \**p* < 0.05; \*\**p* < 0.01; \*\*\**p* < 0.001, \*\*\*\**p* < 0.0001 were considered statistically significant. Tra - trametinib, Veh - vehicle.

Since previous studies have shown that trametinib treatment modulates the TME, thereby enhancing the anti-tumor effect of CD8^+^ T cells (10, 23–27), we stained the vehicle-treated and trametinib-treated tongues with antibodies against the lymphocyte markers CD45 and CD8. IHC analysis of the tongues indicated that the vehicle-treated group was enriched with CD45^+^ cells but the number of CD8^+^ T cells was significantly lower than that in the trametinib-treated group (Fig. 1F). These results suggest that trametinib treatment delays tumor progression by inhibiting the MAPK pathway, reducing tumor cell proliferation, and activating the immune-mediated anti-tumor response by recruiting CD8^+^ T cells.

### CD8^+^ T cells are required for a prolonged response to trametinib

To explore whether the efficacy of trametinib is affected by host immunity, we utilized HNC cell lines derived from 4NQO-induced tumors (28). First, we evaluated the susceptibility of two 4NQO-induced murine HNC cell lines, 4NQO-T (tongue) and 4NQO-L (lip), to trametinib. The half inhibitory concentration (IC_50_) of trametinib was found to be ∼37.5 nM and ∼28.4 nM for 4NQO-T and 4NQO-L, respectively (Fig. 2A). Western blot analysis showed a dose-dependent reduction of pERK levels in both cell lines following trametinib treatment (Fig. 2A). Genomic sequencing of 468 cancer-related genes using the MSK-IMPACT platform (29) showed that these murine cell lines harbor many of the mutated genes found in HNC cancer patients (Supplementary Fig. S2A and Supplementary Table S4). Among these mutated genes, *KRAS* was the only MAPK-pathway related gene that was altered in both cell lines. Specifically, the 4NQO-T cell line harbors a mutation at G12A, while the 4NQO-L line harbors a mutation at G12C; both mutations are present in HNC patients (Supplementary Fig. S2B). To investigate the contribution of the immune system in the response to trametinib, we injected 4NQO-L cells orthotopically into immunocompromised NOD/SCID/IL2rγ^null^ (NSG) and immunocompetent WT mice and compared tumor growth between the mice treated with trametinib and those receiving vehicle alone. In the NSG mice, trametinib treatment slowed tumor progression, while in the WT mice it induced a stable disease with no significant change in the tumor volume over the first 20 days (Fig. 2B and Supplementary Fig. S2C). IHC analysis of the tumors at the end of the experiment showed that while pERK staining was equally reduced in trametinib-treated NSG and WT mice, the proliferation rate of tumor cells was reduced in trametinib-treated WT mice compared to NSG mice (Supplementary Fig. S2D). These results suggest that the host anti-tumor immunity may enhance trametinib efficacy, thereby preventing tumor progression.

**Figure 2.**
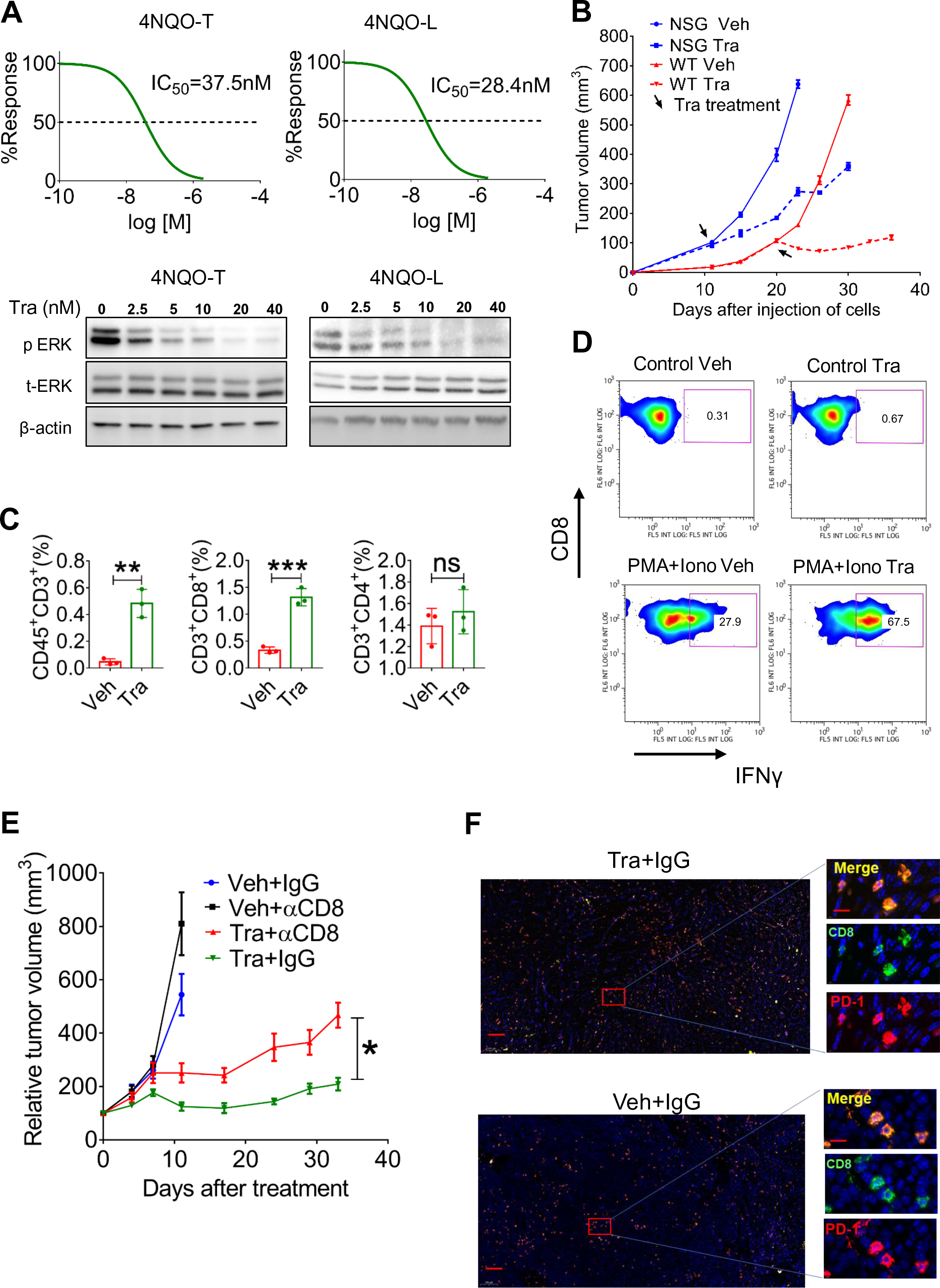
Trametinib treatment induces infiltration of activated CD8^+^ T cells leads to prolonged tumor growth arrest. **(A)** Top – Viability of 4NQO-T and 4NQO-L cell lines treated with increasing doses of trametinib for 4 days; IC_50_ values are shown. Bottom – Western blot analysis showing the expression levels of pERK, total ERK (t-ERK), and beta-actin (as the control) after treatment with increasing doses of trametinib for 24 h in 4NQO-T and 4NQO-L cell lines. (**B)** Growth of 4NQO-L tumors in NSG and WT mice treated with trametinib (Tra) or vehicle. **(C)** Flow cytometry analysis of the lymphocytic population in the tumors treated with vehicle or trametinib for 5 days. **(D)** Intracellular staining of IFNγ in CD8^+^ T cells isolated from the vehicle-treated or trametinib SE-treated (5 days) mice. Density plots showing the percentage of CD8^+^IFNγ^+^ with or without activation with phorbol 12-myristate 13-acetate (PMA) and ionomycin (Iono). Brefeldin A was used as protein transport inhibitor. (**E)** Growth of 4NQO-L tumors in WT mice treated with trametinib, with and without depletion of CD8^+^ T cells. The tumor growth curve is shown for each mouse in each group (n=5–6). (**F)** Immunofluorescence co-staining of CD8^+^ (green) and PD-1 (red) and the merged images (yellow) (scale bars: 200 µm; inset 10 µm). One way ANOVA was performed \**p* < 0.05; \*\**p* < 0.01; \*\*\**p* < 0.001, \*\*\*\**p* < 0.0001 were considered statistically significant. Tra - trametinib, Veh - vehicle.

We then explored whether trametinib affects the infiltration of lymphocytes into the tumors. Analysis of the samples after 5 days of treatment with trametinib revealed a significant increase of CD8^+^ T cells but not CD4 T cells when compared to vehicle-treated tumors (Fig. 2C). Thereafter, we examined the activation status of CD8^+^ T cells in tumors before and after trametinib treatment by comparing their ability to produce interferon-gamma (IFNγ) in response to activation with phorbol 12-myristate 13-acetate (PMA) and ionomycin in vitro(30, 31). Specifically, flow cytometry analysis of PMA and ionomycin-stimulated lymphocytes obtained from tumors of mice treated with vehicle or trametinib for 5 days showed that CD8^+^ T cells derived from trametinib-treated tumors expressed high levels of IFNγ compared to CD8^+^ T cells obtained from vehicle-treated tumors (Fig. 2D and Supplementary Fig. S2E). The enhanced activity of CD8^+^ T cells was also associated with a reduction in the expression of TIM3 (T-cell immunoglobulin domain and mucin domain 3) and PD-1, which are markers of dysfunction and exhaustion of CD8^+^ T cells (Supplementary Fig. S2F)(32–34).

The next step was to investigate the role of CD8^+^ T cells in modulating trametinib efficacy by depleting CD8^+^ cells in 4NQO-L tumor-bearing mice. Specifically, a cohort of WT 4NQO-L tumor-bearing mice was divided into four treatment arms, i.e., vehicle + IgG, trametinib + IgG, vehicle + anti-CD8, and trametinib + anti-CD8. In the mice treated with anti-CD8 (i.e., depleted of CD8^+^ T cells), tumor progression was rapid, compared to the IgG control groups, and in those depleted of CD8^+^ T cells and treated with trametinib the anti-tumor activity of trametinib was attenuated, as evidenced by 4 out of 6 mice developing tumors >300 mm^3^ after 30 days of trametinib treatment. In contrast, at the same time point, none of the tumor-bearing mice treated with trametinib and IgG exhibited tumors larger than 300 mm^3^. Moreover, in mice treated with trametinib + anti-CD8, tumor progression was initiated within 4–5 days, while in the trametinib + IgG group tumor progression started after 20–25 days (Fig. 2E and Supplementary Fig. S2G). Depletion efficiency of CD8^+^ T cells was confirmed by IHC staining for CD8^+^ in the tumors on day 30 (Supplementary Fig. S2H). Co-immunofluorescence (IF) staining of CD8 and PD-1 in vehicle-treated tumors and in tumors that relapsed under trametinib treatment (trametinib + IgG on day 30) showed that CD8^+^ T cells co-expressed PD-1, representing an exhausted phenotype (Fig. 2F). These findings indicate that CD8^+^ T cells are involved in limiting 4NQO-L progression and in promoting the efficacy of trametinib and that prolonged treatment with trametinib creates an immune-suppressive environment, which eventually leads to CD8^+^ T cell dysfunction.

### The trametinib/αPD-1 combination results in tumor elimination and immune memory

Since CD8^+^ T cells showed an exhausted phenotype in the 4NQO-L tumors, and given that MAPK-pathway mutated HNCs are considered to be “hot” tumors (enriched with CD8^+^ cells) and are thus susceptible to αPD-1therapy (4), we predicted that blocking PD-1 would suppress tumor progression of KRAS-mutated 4NQO-T and 4NQO-L tumors. However, αPD-1 monotherapy had no or minimal anti-tumor effects in both 4NQO-L and 4NQO-T HNC models, with some mice showing a tumor growth delay and others being completely resistant to the treatment (Supplementary Fig. S3A). We then posited that further activation of CD8^+^ T cells by supplementing αPD-1 with trametinib would produce superior anti-tumor activity vs. single agents. To explore this premise, we injected 4NQO-L and 4NQO-T cells orthotopically into WT mice. When tumors (3–4 mm diameter) formed, mice were randomized into four groups (vehicle + IgG, vehicle + αPD-1, trametinib + IgG, or trametinib + αPD-1). The 40-day treatment protocol comprised administration of vehicle or trametinib for 5 days, followed by supplementation with IgG or αPD-1 twice a week for an additional 35 days. Analysis of the 4NQO-L tumor volumes and the overall survival of 4NQO-T tumor-bearing mice revealed that treatment with αPD-1 resulted in a transient response that was followed by tumor progression within days. Tumor-bearing mice treated with trametinib exhibited a delay in tumor growth, but within 30 days all mice developed resistance and the tumors progressed (Fig. 3A, B). However, the trametinib/αPD-1 combination induced a profound and durable anti-tumor response in both cancer models. In the 4NQO-L tumor-bearing mice, the combination therapy completely eradicated the tumors in all the mice (6/6) (Fig. 3A). In these animals, tumor relapse was not observed 100 days after completion of the treatment. In the 4NQO-T model, we noted a similar trend with complete elimination of the tumors in 4 out of 6 mice (Fig. 3B). In the 4NQO-T model, disease relapse occurred in 2/6 mice 40 days after the treatment was stopped, suggesting that prolonged treatment with the combined therapy may have prevented tumor relapse.

**Figure 3.**
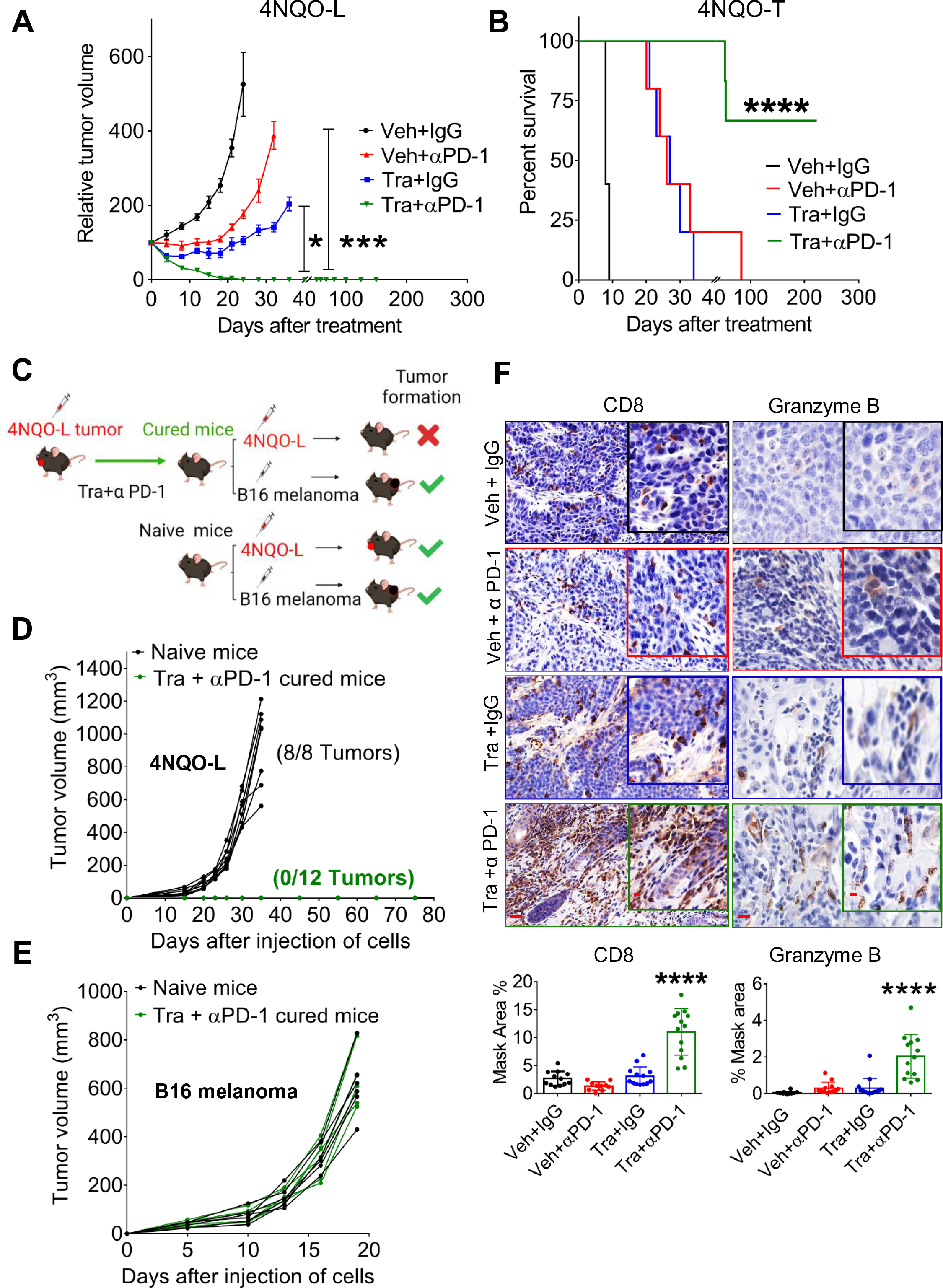
Combination of trametinib and αPD-1 leads to tumor elimination and acquisition of immune memory. **(A)** Volume of 4NQO-L tumors in WT mice treated with αPD1, trametinib or the combination of αPD1 and trametinib. (**B)** Survival of 4NQO-T-tumor bearing WT mice treated with αPD1, trametinib, or a combination of αPD1 and trametinib. (**C)** Scheme showing the re-challenge experimental setting. (**D)** Growth curves of 4NQO-L tumors in naïve and cured mice. (**E)** Growth curves of B16 tumors after injecting naïve and cured mice with B16 melanoma cells. (**F)** Staining and quantification of CD8^+^ T cells and granzyme B in 4NQO-L tumors treated for 31 days as indicated in supplementary fig S3c (Scale bars: 20 µm; insets 10 µm). One way ANOVA was performed, \**p* < 0.05; \*\**p* < 0.01; \*\*\**p* < 0.001, \*\*\*\**p* < 0.0001 were considered statistically significant. Tra - trametinib, Veh - vehicle.

To explore whether the mice in which the combined trametinib/αPD-1 treatment had eliminated the tumors retained long-term immune-memory, we re-challenged the animals by injecting 4NQO-L and B16 melanoma cell lines into the right and left flanks, respectively. A control group of naïve mice was similarly injected with 4NQO-L and B16 melanoma cell lines (Fig. 3C). All naïve mice injected with B16 melanoma and 4NQO-L cells developed measurable tumors. The cured mice rejected the 4NQO-L cells but developed tumors after inoculation with B16 melanoma cells (Fig. 3D, E, and Supplementary Fig. S3B). To confirm the involvement of CD8^+^ T cells in tumor elimination, we repeated the same experiment as that described in Fig. 3A and analyzed some of the tumors before tumor elimination (Supplementary Fig. S3C,D). Histopathological analysis indicated that the tumor shrinkage induced by the trametinib/αPD-1 combination was associated with a massive infiltration and accumulation of activated CD8^+^ T cells, as determined by granzyme B and CD8 staining (Fig. 3F). Taken together, these results indicate that activation of CD8^+^ T cells by the trametinib/αPD-1 combination eliminated KRAS-mutated HNC and that that treated mice had acquired immune memory.

### Chronic treatment with trametinib prevented sensitization of tumors to αPD-1

Given that HNC patients can be treated with compounds targeting the MAPK pathway, such as EGFR inhibitors, prior to immunotherapy prompted us to explore how the duration of pre-treatment with trametinib influences the sensitivity of tumors to supplementation with αPD-1. Specifically, we tested whether a short exposure (SE) or a prolonged exposure (PE) with trametinib determines the ability of αPD-1 to eliminate HNC tumors. The experimental set-up comprised five groups of 4NQO-L-bearing mice (Fig. 4A), treated as follows: vehicle + IgG; vehicle + αPD-1; trametinib + IgG; SE of trametinib for 5 days before starting treatment with αPD-1; and PE of trametinib for 25 days before starting treatment with αPD-1. As expected, supplementation with αPD-1 after a SE with trametinib resulted in tumor eradication in all mice. However, supplementation with αPD-1 after a PE with trametinib arrested tumor growth only for a few days, and the tumors started to relapse thereafter (Fig. 4B). This tumor progression in the trametinib + αPD-1 PE group indicated that trametinib induces a therapeutic window during which αPD-1 efficacy is enhanced, and when mice are exposed to trametinib for an extended period of time, the tumors will regain their αPD-1 resistance phenotype.

**Figure 4.**
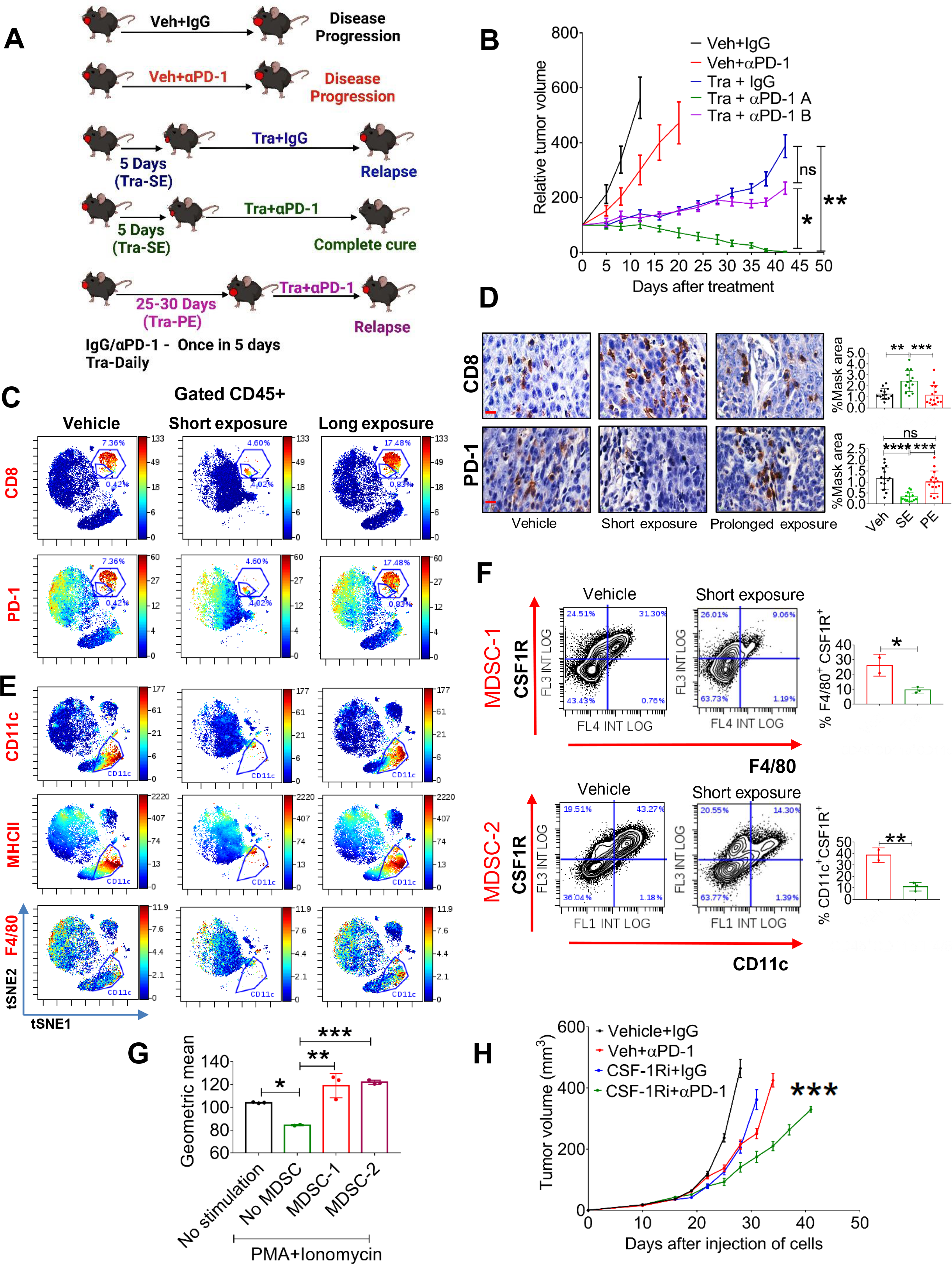
Chronic treatment with trametinib prevents sensitization of tumors to αPD-1. **(A)** Scheme of the experimental setting. (**B)** Tumor volumes of 4NQO-L tumors treated as indicated. (**C)** viSNE plots of the CyTOF data showing CD8 and PD-1 expression in CD45^+^ cells from 4NQO-T tumors treated with vehicle, short exposure (SE; 5 days) trametinib, or a prolonged exposure (PE; 33 days) of trametinib. **(D)** IHC staining (left) of CD8 and PD-1 in 4NQO-L tumors after treatment with vehicle, SE of trametinib, or PE of trametinib (scale bars: 10 µm). Quantification on the right. **(E)** viSNE plots of the CyTOF data showing CD11c, MHCII and F4/80 expression in CD45^+^ cells from 4NQO-T tumors treated with vehicle, SE of trametinib, or PE of trametinib. **(F)** Flow cytometric dot plot analysis of a macrophage-like MDSC population (MDSC-1) and a DC-like MDSC population (MDSC-2) in 4NQO-L tumors treated with vehicle or trametinib for 5 days. (**G)** In-vitro proliferation assay of CD8^+^ T cells in co-culture (ratio 1:10) with two cell populations, MDSC-1 and MDSC-2, derived from 4NQO-L tumor-bearing mice. Geometric mean fluorescent intensities of CFSC (Carboxyfluorescein diacetyl succinimidyl ester) is shown. **(H)** Tumor growth curves of mice injected with 4NQO-L cells in the lip and treated with veh+IgG, CSF-1R inhibitor (PLX-3397), veh+αPD-1, or combination of CSF-1Ri and αPD-1. One way ANOVA was performed, and \**p* < 0.05, \*\**p* < 0.01, \*\*\**p* < 0.001, and \*\*\*\**p* < 0.0001 were considered statistically significant. Tra - trametinib, Veh - vehicle.

To investigate in-depth the effect of trametinib on the heterogeneity of immune cells in the TME of the KRAS-mutated tumors, and to determine whether there is an association between the presence and accumulation of immune cells in the TME and responsiveness to αPD-1, we profiled CD45^+^ cells during trametinib treatment and compared the effect of SE (7 days) vs PE (33 days) of trametinib by using cytometry by time of flight (CyTOF). viSNE analysis of the cytometry data showed that a SE of mice to trametinib resulted in major changes in CD45^+^ cells compared to the vehicle-treated group, while the TME of tumors exposed to trametinib for 33 days had reverted to almost the same landscape as the vehicle-treated group (Fig. 4C). Analysis of CD8^+^ T cells supported the flow cytometry data presented in Fig. 2C and Supplementary Fig. S2F, showing that trametinib treatment induced a reduction of PD-1-positive exhausted CD8^+^ T cells and an increase of activated CD8^+^ T cells (Fig. 4C). Specifically, the percentage of activated CD8^+^ T cells increased from 0.4% in the vehicle-treated tumors to 4% in the SE trametinib-treated tumors, while the exhausted CD8^+^ T cells were reduced from 7% to 4.6%. In contrast, PE with trametinib induced the accumulation of CD8^+^ T cells expressing PD-1 (Fig. 4C). We then quantified by IHC analysis the number and activity of CD8^+^ T cells in both cancer models, 4NQO-L and 4NQO-T, after a SE of 5–10 days or a PE of 25–35 days of treatment with trametinib. Tumors treated with trametinib for 5–10 days showed massive infiltration of CD8^+^ T cells into the tumor site compared to the vehicle-treated group (Fig. 4D and Supplementary Fig. S4A, B). Notably, in the vehicle-treated tumors we observed a larger amount of CD8^+^ T cells at the tumor edge, while in the trametinib-treated group the infiltrated CD8^+^ T cells also spread inside the tumor bulk (Supplementary Fig. S4C). Analysis of the PE group (30 days of treatment) showed a reduction in the number of CD8^+^ T cells inside the tumor, and these CD8^+^ T cells regained their exhausted phenotype with upregulation of PD-1 expression (Fig. 4D and Supplementary Fig. S4A, B).

Analysis of the myeloid compartment showed that CD11b^+^ cells also underwent major alterations after trametinib administration, with elimination of the CD11b^+^CD11c^+^MHCII^+^PD-L1^+^ subpopulation in the trametinib-treated group (Fig. 4E and Supplementary Fig. S4D). The presence of the CD11b^+^CD11c^+^MHCII^+^PD-L1^+^ subset with exhausted CD8^+^ T cells in the TME prompted us to explore whether this CD11b^+^CD11c^+^ population has immunosuppressive capability against CD8^+^ T cells and to determine why a SE with trametinib led to a reduction in the number of these cells. Flow cytometry analysis of the tumors treated with vehicle or trametinib for 5 days showed that the CD11b^+^CD11c^+^MHCII^+^ cells constitute a heterogeneous population expressing CSF-1R^+^ (Supplementary Fig. S4E). One subpopulation of CSF-1R^+^ cells comprises M2-like macrophages that express high levels of F4/80 and MGL2 (CD103b^+^), and the other population shows a monocytic (dendritic cell) DC-like phenotype with F4/80^negative^ and CD11c^+^ (35) cells (Supplementary Fig. S4F, G). Quantification of these subsets after 5 days of treatment with trametinib showed a reduction of the M2-like macrophages, namely CSF-1R^+^ CD11b^+^CD11c^+^F4/80^+^ cells (MDSC-1), and of the monocytic DC-like CSF-1R^+^ CD11c^+^MHCII^+^F4/80^-^ cells (MDSC-2) (Fig. 4F, and Supplementary Fig. S4F). To test whether these myeloid subpopulations have immunosuppressive effects on CD8^+^ proliferation, we co-cultured sorted MDSC-1 (CD11b^+^CD11c^+^MHCII^+^CSF-1R^+^F4/80^+^) and MDSC-2 (CD11b^+^CD11c^+^MHCII^+^ F4/80^-^CSF^-^1R^+^) cells with naïve CD8^+^ T cells and measured CD8+ T cell proliferation in response to stimulation with PMA and ionomycin. The MDSC1 and MDSC2 subsets significantly suppressed the proliferation of PMA/ionomycin-induced naive CD8^+^ T cells (Fig. 4G). In light of the known role of CSF-1R^+^ cells in αPD-1 efficacy(36, 37), we tested whether depletion of CSF-1R^+^ cells in the 4NQO-L tumors would be sufficient to enhance αPD-1 efficacy and to eliminate the tumors. To this end, we treated 4NQO-L tumor-bearing mice with the CSF1R inhibitor PLX3397 (pexidartinib) and supplemented the therapy with αPD-1 or IgG (control). The PLX3397 treatment only partially sensitized tumors to αPD-1 as only a tumor growth delay (and not tumor elimination) was detected in these mice (Fig. 4H). Treatment of mice with PLX3397 was sufficient to reduce the CD11c^+^ population in the tumors, but it did not induce infiltration of CD8^+^ T cells (Supplementary Fig. S4H). Moreover, treatment with the combination of PLX3397 and αPD-1 resulted in a modest (but significant) infiltration of CD8^+^ T cells compared to vehicle which induced only a tumor growth delay and not tumor elimination (Supplementary Fig. S4H).

### Tumor-derived CSF-1 determines CD8^+^ T cell activity and therapy efficacy in mice and in patients

To explore how trametinib treatment reduces the number of CSF-1R^+^ cells and increases the infiltration ability and activation of CD8^+^ T cells, we initially used a publicly available database of single-cell RNA sequencing of HNC patients to examine whether CSF-1, the ligand of CSF-1R, is expressed by tumor cells. Analysis of a cohort of 9 HNC patients showed that some tumor cells, primarily those belonging to the basal and the atypical subsets, express CSF-1 (Supplementary Fig. S5A)(38). We then explored whether trametinib treatment affects the transcriptional levels of CSF-1 in tumor cells. qPCR analysis of 4NQO-L cells treated with trametinib for 12, 24 and 48 h showed decreased CSF-1 levels in vitro (Fig. 5A). However, when 4NQO-L cells were exposed to trametinib for several weeks, no reduction of CSF-1 was detected. We then overexpressed CSF-1 in epithelial 4NQO-L cells and evaluated the tumor response to trametinib in vivo, including profiling the number of MDSCs expressing CD11c^+^ and CD8^+^ T cells in the tumor. The expression level of CSF-1 by the overexpressing cells, 4NQO-L^CSF1^, was three times higher than that of the control cells expressing GFP (4NQO-L^GFP^) (Supplementary Fig. S5B). Analysis of the tumor growth in the control WT mice showed that 4NQO-L^CSF1^ cells grew slightly faster than 4NQO-L^GFP^ cells, but more significant differences in tumor volume were observed after trametinib treatment. While trametinib treatment induced tumor growth arrest for 20 days in 4NQO-L^GFP^ tumor-bearing mice, rapid tumor progression was observed in 4NQO-L^CSF1^ tumor-bearing mice (Fig. 5B). Notably, no differences in the sensitivity of the two cell lines to trametinib treatment were observed in vitro (Supplementary Fig. S5C). IHC analysis of tumors after 5 days of treatment with trametinib showed massive infiltration of CD8^+^ T cells into 4NQO-L^GFP^ tumors, but this infiltration was attenuated in 4NQO-L^CSF1^ tumors, and significantly fewer CD8^+^ T cells were present in trametinib-treated 4NQO-L^CSF1^ tumors compared to 4NQO-L^GFP^ tumors (Fig. 5C, Supplementary Fig. S5D). Moreover, the number of CD11c^+^ cells was higher in 4NQO-L^CSF1^ tumors than in 4NQO-L^GFP^ tumors, while in the 4NQO-L^CSF1^ tumors treated with trametinib the number of CD11c^+^ cells remained as high as the number in 4NQO-L^GFP^ tumors (Fig. 5C, Supplementary Fig. S5D). Based on the negative association between CSF-1 and infiltration of CD8^+^ T cells, we assumed that CSF-1 is a critical factor in regulating the CSF-1R^+^CD11c^+^ MDSC cells in the TME and can thus prevent tumor elimination mediated by the trametinib/αPD-1 combination. To test this hypothesis, we injected 4NQO-L^GFP^ and 4NQO-L^CSF1^ cells into WT mice and compared the efficacy of the trametinib/αPD-1 combination in eliminating tumors in the two groups of mice. Indeed, we observed that combined trametinib/αPD-1 therapy led to the shrinking of the 4NQO-L^GFP^ tumors, and in 5 out of the 6 mice tumors were eliminated within 35-41 days (Fig. 5D and Supplementary Fig. S5E). In contrast, following administration of the combined trametinib/αPD-1 therapy to mice bearing the 4NQO-L^CSF-1^ tumors, 3 out of the 6 tumors progressed, and 3 showed stable disease (Fig. 5D and Supplementary Fig. S5E). These results indicate that CSF-1 served as a key player in modulating CSF-1R^+^CD11c^+^ MDSCs in the TME, thereby preventing the tumor elimination induced by αPD-1 supplementation.

**Figure 5.**
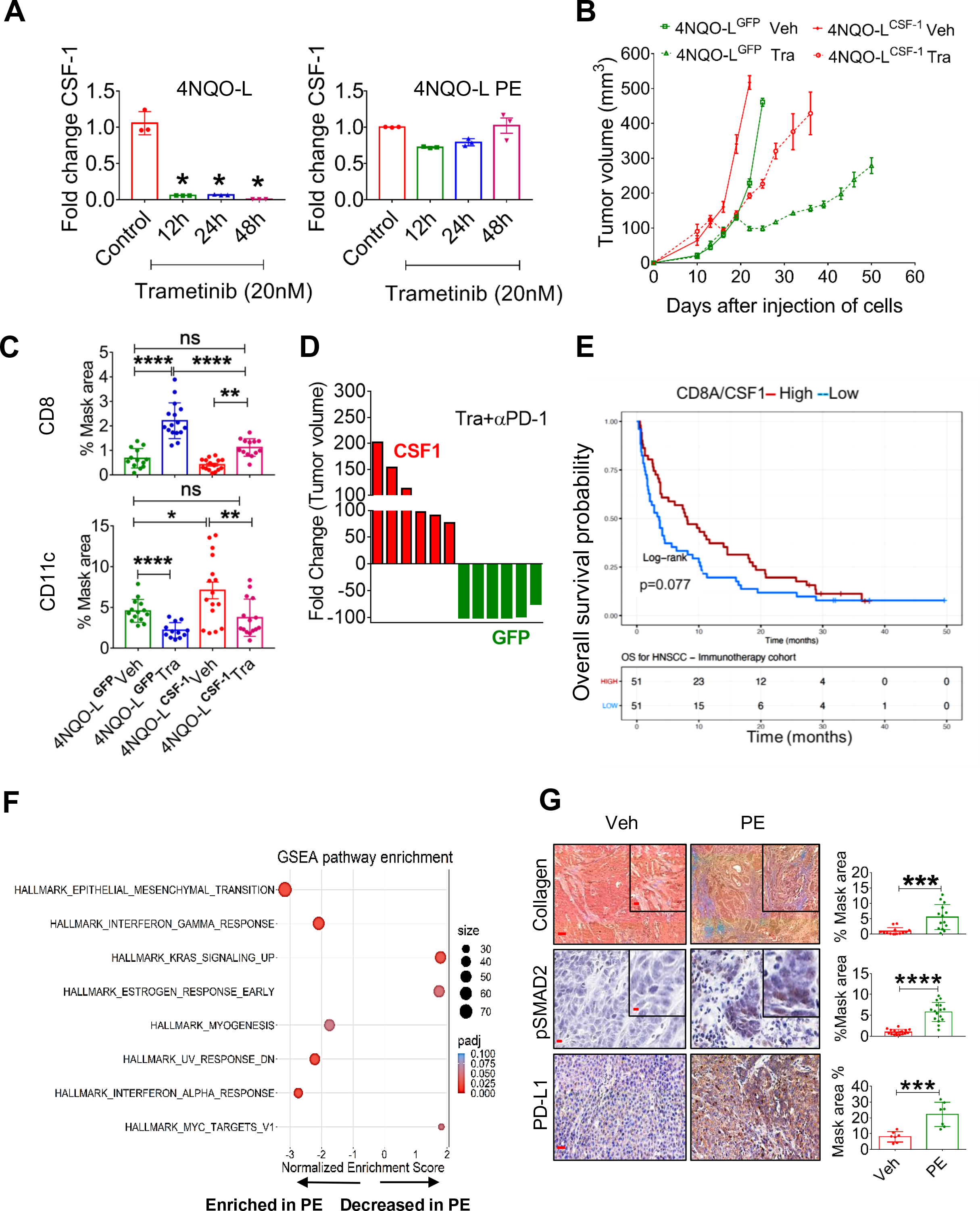
Chronic exposure of trametinib upregulates CSF-1, induces EMT, and prevents trametinib/αPD-1 efficacy. **(A)** mRNA expression levels of CSF-1 in 4NQO-L (left) and 4NQO-L-PE (right) cells after treatment with trametinib (20 nM) for 0, 12, 24 and 48 h. (**B)** Relative volume of 4NQO-L-CSF-1 and 4NQO-L-GFP tumors in WT mice treated with trametinib. (**C)** IHC quantification of CD8^+^ and CD11c in 4NQO-L^GFP^ and 4NQO-L^CSF-1^ tumors treated with trametinib for 5 days (SE). (**D)** 4NQO-L^GFP^ and 4NQO-L^CSF-1^ tumors treated with the combination of trametinib and αPD-1 **(E)** Overall survival (OS) curves for 102 patients with head and neck squamous cell carcinoma (HNSCC) treated with αPD1/PD-L1 (CLB-IHN cohort) and according to high vs. low values of the CD8A/CSF1 ratio. Survival distributions were estimated using the Kaplan-Meier method and compared by the log-rank test between two groups. Patients were binarized at the median. (**F)** GESA analysis of RNAseq of 4NQO-L, 4NQO-L-PE, 4NQO-T, and 4NQO-T-PE cells. **(G)** Collagen (trichrome), pSMAD2 (IHC), and PD-L1 (IHC) in 4NQO-L tumors treated with vehicle or PE of trametinib (scale bars: 100 µm (inset 20 µm), 50 µm (inset 10 µm), and 100 µm respectively). For statistics, an unpaired 2-sided t-test or one-way ANOVA was performed. \**p* < 0.05; \*\**p* < 0.01; \*\*\**p* < 0.001, \*\*\*\**p* < 0.0001 were considered statistically significant. Tra - trametinib, Veh - vehicle.

To explore whether these findings can serve as indicators of response to immunotherapy, we undertook gene expression profiling of 102 confirmed recurrent HNC patients pre-treated with αPD-1/αPD-L1 before the initiation of immunotherapy at Centre Leon Berard (CLB, Lyon France) (Supplementary Table S5)40. By calculating the ratio of the expression of CD8A to the expression of CSF-1, we found that a high CD8A/CSF1 ratio in pretreated patients led to a higher overall survival (binarization at median; p-value = 0.077) with Hazard Ratio of 0.69; 95%CI [0.46;1.04] (Fig. 5E). Consistently, a high CD8A/CSF1 ratio also tended to be associated with a clinical benefit in response to immunotherapy [defined as complete response, partial response, and stable disease for at least six months; p = 0.077) (Supplementary Fig. S5F)]. A similar trend was observed in lung cancer patients, as responders to immunotherapy expressed higher ratio levels of CD8A/CSF-1 than non-responders (p = 0.052) (Supplementary Fig. S5G)(39).

To seek an explanation for the constancy of CSF-1 levels in cells that were chronically exposed to trametinib (Fig 5A), we compared the whole transcriptome of 4NQO-L and 4NQO-T cells before and after a PE to trametinib (Supplementary Table S6). While we did not observe differences in baseline levels of CSF-1, upregulation of genes of the EMT signature were significantly enriched after PE to trametinib (Fig. 5F). Specifically, EMT-associated protein expression, such as VIM, TGFβ2, COL1A1, and PD-L1, was upregulated (Supplementary Table S7). IF and western blot analysis confirmed an EMT shift in cellular plasticity and upregulation of PD-L1 in vitro (Fig. S5H, I). An in vivo comparison of 4NQO-T and 4NQO-L tumors treated with vehicle or trametinib for 25–30 days (PE) showed an increase in PD-L1, pSMAD2, and collagen expression within the TME in the trametinib-treated tumors compared to the vehicle-treated group (Fig. 5G and Supplementary Fig. S5J). We confirmed the association of expression of the EMT genes, *VIM* and *ZEB1*, with CSF-1 in the TCGA dataset of HNC cancer patients (Supplementary Fig. S5K). We then explored the transcription factors (TFs) that may maintain CSF-1 expression in 4NQO-PE cells. To this end, we used the oPOSSUM.3 system(40) to perform TF analysis for the genes upregulated in PE cells (FC<-1, Padj<0.001) and found that 65 TFs were activated (Z-score> 2) (Supplementary Table S8). Cross-section analysis of these TFs with a list of TFs known to bind the CSF-1 promoter extracted from the ENCODE Chip-seq (Supplementary Fig. S5L) showed that AP1, STAT1/3, CTCF, and TBP might be implicated in CSF-1 expression (Supplementary Fig. S5L). Taken together, our findings show that prolonged treatment of KRAS-mutated HNC with trametinib resulted in, signaling and transcriptional adaptation, leading to EMT and maintained CSF-1 levels in the presence of trametinib, thereby prevents the efficacy of supplementation with αPD1.

## Discussion

In this study, we demonstrated that trametinib treatment delayed the initiation and progression of MAPK-pathway-mutated HNC and changed the heterogeneity of the immune cells in the TME, which subsequently determined the susceptibility to trametinib/αPD1 combination therapy. In the immediate response to a short treatment with trametinib, defined as a SE of ∼5–10 days, tumors were enriched with activated CD8^+^ T cells. However, after prolonged treatment (PE of ∼>25 days), tumors adapted to the therapeutic stress and re-established an immunosuppressive environment enriched with exhausted CD8^+^ T cells. Importantly, activation of CD8^+^ T cells with αPD-1 after a SE of trametinib resulted in tumor elimination and the establishment of immune memory, while supplementation with αPD-1 after a PE of trametinib resulted in tumor progression (Scheme 1).

**Scheme 1.**
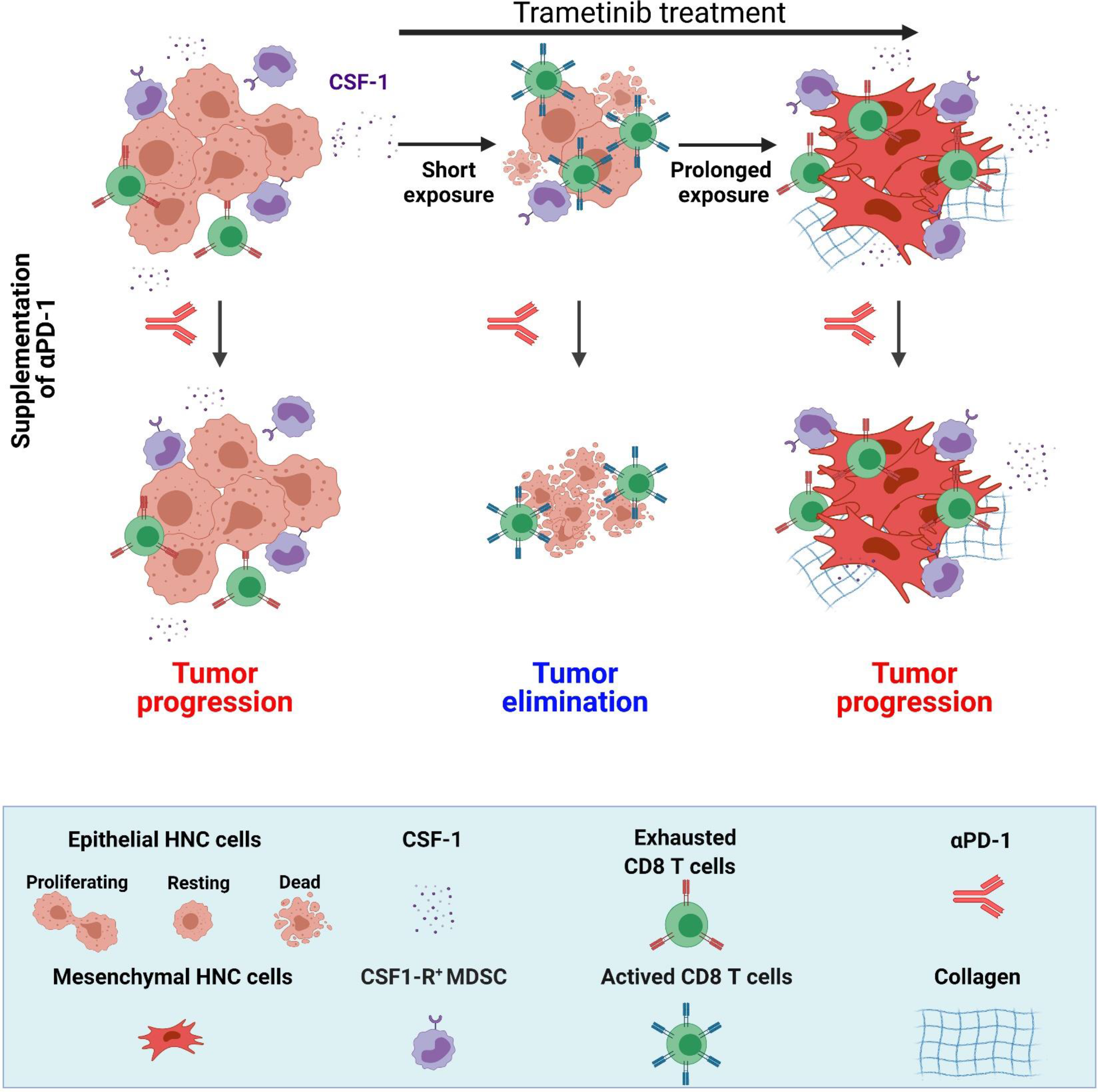
The effect of the duration of trametinib treatment on sensitization of MAPK-pathway mutated HNC to supplementation of αPD-1. MAPK-pathway mutated HNC are resistant αPD-1 and sensitive to trametinib. SE of trametinib leads to the reduction of CSF-1 secretion, reduction of CSF-1R^+^ MDSCs and infiltration of active CD8^+^ T cells while prolonged exposure of trametinib induced EMT phenotype and restored the CSF-1 secretion, CSF-1R^+^ MDSCs and exhaustion of CD8^+^ T cells. Supplementation of αPD-1 after SE of trametinib leads to complete elimination of tumor but supplementation of αPD-1 after PE of trametinib resulted in tumor progression. (Scheme was created with BioRender.com).

Several studies in pre-clinical models and in cancer patients have demonstrated that targeting the MAPK pathway (i.e., with RAF and MEK inhibitors) induces CD8^+^ T cell infiltration into tumors and improved immunotherapy efficacy (25, 41–44). Mechanistically, MAPK pathway inhibition has a dual effect: on the one hand, it downregulates the expression of immunosuppressive factors such as IL-8 and IL-1 and increases the infiltration of activated CD8^+^ T cells, but on the other hand, it counteracts immune activation through upregulation of PD-L1 by tumor cells. Similarly, in our KRAS-mutated HNC tumors, trametinib treatment initially induced temporary immune activation with massive CD8^+^ T cell infiltration and reduced the numbers of CSF1R^+^CD11c^+^ MDSCs, but chronic treatment resulted in immune suppression. The reduction in the accumulation of the MDSCs expressing CSF1R^+^CD11c^+^ seems to be mediated, at least in part, by a reduction in tumor-derived CSF-1. Indeed, ectopic CSF-1 overexpression by tumor cells attenuated the trametinib-induced reduction of CSF1R^+^CD11c^+^ MDSCs and the infiltration of activated CD8^+^ T cells into the TME. CSF-1 is a key pro-tumorigenic chemokine that promotes the accumulation, differentiation, and proliferation of subsets of CSF1R^+^ MDSCs that suppress the activation and proliferation of T cells (45–47). Several reports have shown that high expression of CSF-1 is associated with the suppression of T cells and with resistance to immunotherapies, such as αPD-1 (36, 37, 48). In line with these observations, depletion of CSF-1R^+^ cells with small molecule inhibitors attenuates tumor progression(49) and enhances therapy efficacy(36, 50). However, while the suppressive activity of CSF1R^+^ cells towards CD8^+^ T cells and natural killer cells has been reported in multiple cancers(51–54), targeting CSF-1R^+^ cells in mice bearing 4NQO-L tumors had only a minor effect on tumor growth.

Allegrezza et al. (27) have shown that trametinib reduces MDSC myelopoiesis and exhibits enhanced efficacy against KRAS-driven breast cancer due to the activation of CD8^+^ T cells. Our results are in line with these findings, as trametinib was more potent in immunocompetent (WT) than in immunocompromised (NSG) mice, and depletion of CD8^+^ T cells attenuated the efficacy of trametinib in WT mice. Using 4NQO-L^CSF1^ and 4NQO-L^GFP^ tumors we provided further evidence to support the influence of tumor-derived CSF-1 on CSF1R^+^CD11c^+^ MDSCs and CD8^+^ T cells in response to therapy in vivo. We showed an association between trametinib-induced down-regulation of CSF1 expression and a reduction in CD11c^+^CSF1R^+^ MDSCs with the sensitivity of the tumors to αPD-1. However, depletion of CSF1R^+^CD11c^+^ cells with the CSF-1R inhibitor PLX3397 in combination with αPD1 only induced a tumor growth delay and was not able to eliminate the tumors. These results differ from reports showing that CSF1R^+^ elimination can enhance the efficacy of immunotherapy and cause tumor elimination in other cancer types(48, 54). The minor effect of PLX3397 compared to trametinib in sensitizing tumors to αPD-1 highlights the fact that downregulating CSF-1 is only a part of the pleiotropic role played by trametinib in anti-cancer immunity. This notion is reinforced by the evidence of trametinib-induced growth arrest of KRAS-mutated tumors, accompanied by CD8^+^ T cell infiltration into the tumors and by reprogramming CD8^+^ T cells into memory stem cells with potent anti-tumor effects (20, 26, 27).

One of the key findings of our study is the demonstration of a therapeutic vulnerability induced by trametinib treatment that enables αPD-1 to facilitate HNC eradication in mice. The induction of immune activation following treatments with targeted therapy, chemotherapy, or irradiation to enhance immunotherapy efficacy has been described in HNC patients (55). Recently, Choi et al. showed that pulsatile treatment of KRAS-mutated tumor-bearing mice with selumetinib or trametinib had a greater effect on immune activation and susceptibility to combination therapy with CTLA4 as compared to continuous treatment (26). These results further support the importance of induction by anti-MEK1/2 therapies of a potent and prolonged immune activation.

In our study, as well as in others, the immune activation mediated by trametinib is associated with a delay of tumor growth and, when therapy is subsequently initiated, it is also associated with immune suppression (23, 56). Explanations for our findings may be drawn from previous reports showing that prolonged treatment with trametinib induces signaling adaptation in HNC cells related to the mesenchymal phenotype, which may be promoted by the TF, YAP (57, 58). It is also known that tumors showing an EMT phenotype are enriched with immunosuppression characteristics (59, 60). For example, tumor cells undergoing EMT upregulate immunomodulatory surface proteins, such as PD-L1, and express high levels of chemokines that regulate immune suppressive factors, such as IL-10, TGF-β and CSF-1(61, 62). Thus, a potential explanation for the constancy of CSF-1 expression in PE tumor cells is the hyperactivation of TFs in EMT-like cells, such as TWIST-1, which has been shown to maintain CSF-1 expression (63). Another mechanism that further impairs αPD-1 efficacy in tumors undergoing EMT is the upregulation of collagen, which has been shown to induce CD8^+^ T cell exhaustion and to promote resistance to αPD-1/PD-L1 (64, 65).

Recently Hass et al.(66) demonstrated that the acquisition of resistance to targeted anti-MAPK therapy (with MEK or RAF inhibitors) conferred cross-resistance to immunotherapy in melanoma and they proposed that immunotherapy should be administrated before patients develop resistance to anti-MEK1/2. Their basic observation that prolonged treatment with anti-MEK1/2 resulted in resistance to αPD-1 (also shown in our HNC models) is suggestive of a general phenotype of MEK inhibitors in MAPK-pathway mutated tumors. Nevertheless, the mechanisms may be different in the two sets of experiments: while Hass et al. showed that treatment of tumor-bearing mice with anti-MAPK increased the number of CD103^+^ DC cells in the TME, thereby activating CD8^+^ T cells, we proposed an alternative mechanism based on the elimination of the immune-suppressive CD11c^+^/CSF-1R^+^ cells.

HNCs are classified into four major subtypes based on RNA sequencing (38, 67). MAPK-pathway mutated tumors are associated with the immune subtype that is enriched with CD8^+^ T and CD11c cells(4), while tumors that have hyperactivated MAPK pathway via RTK stimulation are associated with the basal subtype(3). The IHC and CyTOF analyses of the TME of the KRAS-mutated 4NQO-T and 4NQO-L tumors showed a high amount of CD45^+^ cells and an association between CSF1R^+^CD11c^+^ MDSCs and exhausted CD8^+^ T cells, but despite this “immune” enriched phenotype, the tumors were resistant to αPD-1. This resistance to αPD-1 can be explained by the enrichment of dysfunctional/exhausted CD8^+^ T cells with high PD-1 expression, as observed in HNC patients (68), and in 4NQO tumors before and after the PE with trametinib. Our observation that tumors became sensitive to αPD-1 only when trametinib reduced CSF1 expression and induced CD8^+^ infiltration encouraged us to explore whether the ratio between CSF1 and CD8A is associated with a clinical benefit derived from administering αPD1 to HNC patients. An analysis of gene expression profiles from pre-immunotherapy biopsies of HNC patients showed that patients expressing low levels of CSF1 and high levels of CD8A displayed a better overall survival and an association with a clinical benefit of αPD-1 and αPD-L1 therapies.

Although our study was confined in KRAS mutated HNC models, we predict that our phenomenon are not restricted to KRAS mutated tumors as we recently found that KRAS pathway activity might be a common principle for HNC (69). Nonetheless, since it is possible that the results of these studies and those of our study could be selective for KRAS-mutated cancers, further studies are needed to exclude this possibility, which, in turn, underscores the importance of developing new pre-clinical HNC models harboring a wide range of oncogenic mutations.

In conclusion, our findings show the potential of of sensitizing MAPK-pathway mutated HNC to αPD-1 by pre-treating patients with MAPK inhibitors, and then continuing with the therapy combination. Pre-treatment with trametinib interfered with the interaction between the malignant cells and the microenvironment via reducing tumor-derived CSF-1. Downregulation of CSF1 expression and arrest of tumor cell proliferation resulted in an immune-active TME, which is required for immunotherapy efficacy. Such an active immune state of the TME defined by the CD8A to CSF-1 expression ratio is associated with a positive response to αPD1/αPD-L1 therapies in cancer patients.

## Materials and methods

### Genomic analysis of MAPK pathway mutated genes

The MAPK pathway genes were defined in the CBbioPortal (70–73). Whole-exome sequencing data for MAPK pathway-specific mutation was retrieved from TCGA, GENIE, and ICGC for mutational analysis on 9 February 2021 (www.cbioportal.org, www.genie.cbioportal.org, www.dcc.icgc.org).

### MAPK pathway activation - TCGA analysis

A heatmap was generated with the web tool ClustVis(74) MAPK pathway activity was inferred using the progeny R package(75), as described previously (76). GSVA scores were computed with the GSVA package (76) in RStudio (4.0.2) based on selected gene sets from the Molecular Signatures Database (MSigDB (77, 78)).

### Oral carcinogenesis with 4-nitroquinoline N-oxide (4NQO)

C57BL/6J mice, 4–5 weeks old, purchased from the Envigo laboratory, were used for the 4NQO carcinogenesis studies. A stock solution of 4NQO (N8141 Sigma, St. Louis, MO) in propylene glycol was prepared weekly at a concentration of 5 mg/mL and administered at a concentration of 50 μg/mL in the drinking water for 12 weeks. The development of tumors in the oral cavity of the mice was examined weekly, and at end of the experiment, tumors were collected for analysis. For the survival experiments involving trametinib (GSK1120212, JTP-74057), the drug was given after 12 weeks of 4NQO treatment (trametinib was dissolved in 4% DMSO and then diluted in corn oil and administered at a dose of 1 mg/kg/day).

### In vivo experiments

Mice were housed in air-filtered laminar flow cabinets with a 12-h light/dark cycle and supplied with food and water ad libitum. All animal experiments were carried out under the Institutional Animal Care and Use Committee (IACUC) of Ben-Gurion University of the Negev (BGU’s IACUC) according to specified protocols to ensure animal welfare and to reduce suffering. The animal ethical clearance protocol numbers used for the study are IL-80-12-2015 (E), and IL-29-05-2018 (E). In vivo experiments were conducted using 6- to 8-week-old NSG mice (NOD.Cg-Prkdcscid Il2rgtm1Wjl/SzJ, Jackson Labs) and WT mice (Envigo, Huntingdon, UK, C57/BL/6).

### Patient-derived xenografts (PDXs)

For this study, we used two PDXs, designated SE19 (PDX-1) and SE103 (PDX-2), previously established in our lab (79, 80). The tumors were transplanted into the right and left flanks of NSG mice for the trametinib (1 mg/kg/day) efficacy experiments. In all experiments, tumor measurement was performed with a digital caliper, and tumor volumes were determined according to the formula: length × width^2^ × π/6.

### Immunohistochemistry

Tissues were fixed in a 4% paraformaldehyde (PFA) solution for a maximum of 24 h at room temperature, dehydrated, and embedded in paraffin. The tissue sections were de-paraffinized with xylene. H_2_O_2_, 3%, was used to block the endogenous peroxidase activity for 20 min, and thereafter the sections were rinsed in water for 5 min. Antigen retrieval was performed in citrate buffer (pH 6) at 99.99 °C for 15 min. Sections were then blocked for 1 h at room temperature with blocking solution [phosphate buffered saline (PBS), 0.1% Tween (0.0125% for AXL staining), 5% bovine serum albumin (BSA)], followed by incubation with primary antibody (diluted in blocking solution) overnight at 4 °C. A list of antibodies used is given in Supplementary Table S9. The ABC kit (VECTASTAIN Cat. VE-PK-6200) was used for color detection according to the manufacturer’s protocol. Sections were counter-stained with hematoxylin and mounted in mounting medium (Micromount, Leica Cat. 380-1730).

### Cell lines

4NQO-L and 4NQO-T murine cell lines developed as described previously(28) were maintained at 37 °C in a humidified atmosphere at 5% CO_2_ in DMEM supplemented with 1% L-glutamine 200 mM, 100 units each of penicillin and streptomycin, and 10% fetal bovine serum (FBS). Cells were routinely tested for mycoplasma infection and treated with appropriate antibiotics if needed (De-Plasma, TOKU-E, D022). For generating 4NQO-L-PE and 4NQO-T-PE cells, 4NQO-L and 4NQO-T were exposed to 50–100 nM of trametinib for several weeks.

### IC_50_ assay

Cells were seeded in 96-well plates (5000 cells/well), treated with increasing concentrations of trametinib, and allowed to proliferate for 72 h. Cells were then stained with crystal violet (1 g/L) for 10 min, rinsed, and dried, and bound crystal violet was dissolved out with 10% acetic acid. Absorbance was measured at 570 nm (BioTek™ Epoch™ spectrophotometer). The dose-response curve was plotted, and IC_50_ values were calculated using GraphPadPrism7 software.

### Western blotting

Cells were harvested and lysed using lysis buffer (50 mM HEPES, pH 7.5, 150 mM NaCl, 1 mM EDTA, 1 mM EGTA, 10% glycerol, 1% Triton X-100, 10 µM MgCl_2_), supplemented with phosphatase inhibitor (Bio tool, B15001A/B) and protease inhibitor (Millipore Sigma, P2714-1BTL) cocktails, and placed on ice for 30 min, followed by 3 min of sonication. Lysates were cleared by centrifugation (30 min, 14,000 rpm, 4 °C). Supernatants were collected, and whole-cell lysates (25 µg) were separated on 10% SDS–PAGE and blotted onto PVDF membranes (BioRad Trans-Blot® Turbo™ transfer pack #1704157). Membranes were blocked for 1 h in blocking solution [5% BSA (Amresco 0332-TAM) in Tris-buffered saline (TBS) with 0.1% Tween] and then incubated with primary antibodies diluted in blocking solution. Mouse and rabbit horseradish peroxidase (HRP)-conjugated secondary antibodies were diluted in blocking solution. Protein-antibody complexes were detected by chemiluminescence [Westar Supernova (Cyanagen Cat. XLS3.0100) and Westar Nova 2.0 (Cyanagen Cat. XLS071.0250)], and images were captured using the Azure C300 Chemiluminescent Western Blot Imaging System (Azure Biosystems). Details of antibodies used are presented in Supplementary Table S9.

### H&E staining

Rehydrated tissues were stained with hematoxylin for 3 min, washed with tap water for removing excess stain, and stained with eosin for 30 s. Excess stain was removed by dipping in 70% ethanol. Tissues were then dehydrated and mounted.

### Immunofluorescence

For IF monitoring, cells were seeded on 24-mm round coverslips. Cells were washed with PBS and fixed in 4% PFA for 30 min at room temperature. Cells were rinsed with PBS, followed by permeabilization using 0.05% Triton X-100 (Millipore Sigma) for 10 min in PBS. Antigen retrieval for tissue section staining was performed in citric acid. Then, cells/tissues were blocked using a blocking solution (5% BSA in TBS-T) for 1 h at room temperature. Cells/tissues were incubated with primary antibodies overnight at 4 °C, washed in PBS-T and stained with secondary antibodies. Slides were mounted with DAPI Fluoromount-G® (SouthernBiotech, Birmingham, MA, USA, 0100-20).

### Masson’s Trichrome Staining

Tissues were rehydrated and stained using Masson’s Trichome Staining Kit as per the manufacturer’s instructions (Bio-optica-04-01802). Slides were mounted and scanned using the Pannoramic Scanner (3DHISTECH, Budapest, Hungary) and analyzed with a QuantCenter (3DHISTECH, Budapest, Hungary).

### OPAL multiplexed staining of tissues

Tumor tissues were prepared for staining as described above, and then IF was performed as described in the manufacturer’s protocol (Opal™ 4-Color Manual IHC Kit, cat no. NEL810001KT). Briefly, tissues were incubated with an anti-CD8 antibody (Cell Signaling Technology, 1:500) for 1 h at room temperature. Tissues were then washed with Tween-PBS and incubated with rabbit horseradish peroxidase (HRP)-conjugated secondary antibody (Jackson ImmunoResearch, PA, USA, 1:500) for 1 h at room temperature. Thereafter, tissues were again washed and stained with FP1487001KT [Opal™ 570 FP1488001KT (red)], according to the manufacturer’s protocol. The method is based on stripping the primary and secondary antibodies (not reducing fluorescent signals) and then restaining with a second primary antibody, using the same procedure. After stripping, the tissues were incubated with α-PD-1 antibody (Cell Signaling Technology, 1:250) for 1 h at room temperature, washed, and stained again with rabbit HRP-conjugated secondary antibody, followed by staining with FP1487001KT. Slides were mounted with DAPI Fluoromount-G^®^ (SouthernBiotech, Birmingham, MA, USA, 0100-20). IHC and IF slides were scanned using the Pannoramic Scanner (3DHISTECH, Budapest, Hungary) and analyzed with a QuantCenter (3DHISTECH, Budapest, Hungary) using a single threshold parameter for all images with a specific staining in each experiment.

### Orthotopic models for drug efficacy/survival studies

Orthotopic models were established by injecting tumor cell lines into the lips or tongues of the mice. For the efficacy studies, treatments were started when the tumors in the lip had reached 4–5 mm in diameter. Tumors were measured with a digital caliper twice a week. At the end of the experiment, animals were sacrificed, and the tumors were collected. For survival experiments, cells were injected orthotopically into the tongues of the mice. For efficacy experiments with trametinib and αPD-1 antibody, trametinib was used at 1 mg/kg/day, and αPD-1 (rat anti-mouse PD-1, BE0146-25 -clone RMP1-14) and IgG (rat anti-mouse IgG2a, BE0089-25) from Bio X Cell were used at a concentration of 100 µg/mouse. For the efficacy experiments with the CSF-1R inhibitor, pexidartinib (Chemscene #4256), the compound was used at a concentration of 75 mg/kg dissolved in DMSO (4%), and corn oil was used as the vehicle.

### CD8 depletion experiment

In vivo Plus™ anti-mouse CD8α (rat anti-mouse CD8α, BP0061-25 Clone 2.43) or IgG In vivo Plus ™ rat IgG2b isotype control, both from Bio X Cell, were used for the CD8 depletion experiments. The animals were given IP, 1 mg/mouse of αCD8 antibody or IgG before 2 days of trametinib treatment, and treatment was continued with 500 µg/mouse of αCD8 antibody or IgG every 5 days.

### CyTOF

#### Sample preparation

The tumor tissue was washed with PBS, cut into small pieces, and suspended in 1 mg/mL type II collagenase diluted in RPMI 1640 medium. The gentleMACS™ Octo Dissociator (Miltenyi Biotec, Bergisch Gladbach, Germany) was used to dissociate the tumor tissue at 37 °C for 35 min. Then, the samples were filtered through a 40-μm strainer to obtain a single-cell suspension. Erythrocytes were removed with ACK lysing buffer [0.15 M NH4Cl, 10 mM KHCO3, and 0.5 M EDTA (pH 7.2–7.4)]. Cells were then washed once with cell staining medium (CSM; PBS, 2% BSA and 0.07% azide), and samples were resuspended in 500 μL of Cell-ID Intercalator-103Rh (1:2000) (Fluidigm) for 15 min at room temperature or overnight at 4 °C. The cells were washed in CSM and resuspended in 50 μL of CSM. Most antibodies were obtained pre-conjugated to heavy-metal isotopes from Fluidigm. Cell-surface antibody master mix was prepared by adding appropriate dilutions of all cell-surface antibodies into 50 μL of CSM per sample. The antibody master-mix was then filtered through a pre-wetted 0.1-μm spin-column (Millipore) to remove antibody aggregates, and 50 μL of this filtrate were added to the sample resuspended in 50 μl of CSM. After incubation for 60 min at room temperature, cells were washed once with CSM and fixed in 500 μL of 1.6% PFA in PBS and stored at 4 °C. Cells were washed once with CSM and resuspended in intercalation solution [1.6% 0.5 μM iridium-intercalator (Fluidigm)] for 20 min at room temperature or overnight at 4 °C.

#### Data acquisition

Before data acquisition, samples were washed once in CSM and twice in doubly distilled water and filtered through a cell strainer (Falcon). Cells were then resuspended at 5 × 10^6^ cells/mL in doubly distilled water supplemented with 1× EQ four-element calibration beads (Fluidigm), and data was acquired on a CyTOF2 mass cytometer (Fluidigm). The data was then bead-normalized using MATLAB-based software(81). The normalized data was uploaded onto the Cytobank analysis platform(82) to perform initial gating and population identification using the indicated gating schemes. Data was represented as viSNE plots. viSNE schemes were created using the Cytobank online tool.

### Flow cytometry

The gentleMACS™ Octo Dissociator (Miltenyi Biotec, Bergisch Gladbach, Germany) was used to dissociate the tumor tissue at 37 °C for 35 min.. Samples were filtered through a 40-μm cell strainer to obtain a single-cell suspension. Cells (1 million/sample) were stained following blocking with anti-CD16/32 (anti-FcγIII/II receptor, clone 2.4G2, 20 min) with the primary antibodies for 20–30 min on ice. DAPI/Aqua dead cells marker was used to gate the dead cells. Details of the fluorescent-labeled antibodies are listed in Supplementary Table S9. Samples were analyzed in a Gallios flow cytometer. Data were analyzed using FlowJo software or the Cytobank online tool.

### Intracellular cytokine staining

CD8^+^ T cells were isolated from tumors treated with vehicle or trametinib for 5 days and activated with 25 ng/mL PMA [Santa Cruz Biotechnology (sc-3576)] and 1 µM ionomycin [Santa Cruz Biotechnology (sc-3576)] for 5 h. After activation, cells were stained with the appropriate surface markers [dead cell marker-Zombie Aqua™ Fixable Viability Kit (Biolegend)] and CD8 (APC conjugated) and then fixed using 4% PFA for 20 min at room temperature. Then cells were permeabilized with permeabilization buffer (1X) (eBioscience -00-8333-56) and stained with IFNγ-PE-cy7 for 20 min. Cells were then washed, and flow cytometry analysis was performed using Gallios flow cytometer. Data were analyzed using FlowJo software.

### In vitro T-cell proliferation and activation assays

MDSCs, i.e., CD11b^+^, F4/80^+^, CD11c^+^, MHC11^+^ CSFR^+^, CD11b^+^, F480^-^, CD11c^+^, and MHC11^+^ CSFR^+^ cells, were sorted from tumors using a FACSAria II flow cytometer. Naive CD8+^+^ T cells were purified from naïve C57/BL6 mice by using the mouse CD8^+^ T Cell Isolation Kit (Miltenyi, 130-104-075). MDSCs and CD8^+^ T cells were co-cultured at a ratio of 1:10 for 96 h. T cells were activated with 25 ng/mL PMA and 1 µM ionomycin. Proliferation of CD8^+^ T cells was determined using the CFSE Cell Division Tracker Kit (Biolegend 423801) according to the manufacturer’s protocols.

### Generation of CSF-1-overexpressing cell lines

A lentiviral vector for CSF-1 (pLV-EGFP:T2A:Puro-EF1A>mCSF-1 [NM_001113530.1] and an EGFP lentiviral control vector (pLV-EGFP/Puro-CMV>Stuffer300) were ordered from VectorBuilder. 4NQ-O cells were infected with lentiviruses encoding for murine CSF-1 and EGFP as control, and the infected cells were selected using puromycin (5µg/mL).

### RNA sequencing

RNA sequencing was performed on the 4NQO-L, 4NQO-T, 4NQO-L-PE, and 4NQO-T-PE cells. According to the manufacturer’s instructions, RNA was extracted using the RNeasy mini kit (Qiagen, 74104). RNA-seq libraries were prepared as described previously by Elkabets lab(79). Briefly, RNA-seq libraries were prepared using the Tru Seq RNA Sample Preparation kit (Illumina, San Diego, CA, USA) according to the manufacturer’s protocol. Total RNA, 1 mg, was fragmented, followed by reverse transcription and second-strand cDNA synthesis. The DS-cDNA was subjected to end repair, a base addition, adapter ligation, and PCR amplification to create libraries. Sequencing was performed with a Nextseq 5000 system, using all four lanes.

### Real-time quantitative PCR

Total RNA was isolated from 4NQO-L, 4NQO-T, 4NQO-L-PE, and 4NQO-T-PE cells treated with trametinib (20 nM) for different times, using ISOLATE II RNA Mini Kit (Bioline-BIO-52073) according to the manufacturer’s protocol. RNA, 1 μg, was converted to cDNA using a qScript cDNA synthesis kit (Quanta Bioscience, 95047-100) according to the manufacturer’s protocol. Real-time PCR was performed (Roche LightCycler® 480 II) using a prime time gene expression master mix (IDT, 1055770), with matching probes from IDT: CSF-1 gene mouse (Mm.PT.58.11661276) and GAPDH mouse (Mm.PT.39a.1).

### CLB-IHN cohorts

The CLB-IHN cohort is derived from a previously published cohort of patients treated at CLB (Lyon France) for a histologically confirmed recurrent or metastatic head and neck SCC in clinical trials testing the efficacy of PD-1/PD-L1 antibodies alone or in combination with an anti-KIR or an anti-CTLA4 antibody between March 2014 and November 2018(83) (Supplementary Table S5). Targeted gene expression profiles (HTG EdgeSeq technology, Oncology Biomarker Panel [OBP]) were generated for pre-immunotherapy formalin-fixed, paraffin-embedded (FFPE) tumor samples. Briefly, the median age of the patients was 63 years (range 33-88). Most patients were male (81%), current/former smokers (87%), and alcohol drinkers (85%). HNC originated mainly in the oropharynx (39%) and oral cavity (33%) and best response on immune checkpoint immunotherapy were only 11% with partial response and complete response.

### Statistical analysis

Each experiment was repeated 2 or 3 times, and representative data are shown. Statistical analysis was performed using GraphPad Prism 7 software, and results are presented as means ± SEM. IHC images were analyzed by Histoquant software (3DHISTECH), and differences between the two groups were analyzed using an independent t-test, whereas for the analysis of more than two groups a one-way ANOVA test was used. Correlation analysis was conducted using Spearman’s rho test. Overall survival (OS) was defined by the time in months from tumor biopsy to death or loss to follow-up. In the CLB-IHN cohort, survival distributions were estimated using the Kaplan–Meier method and compared with the log-rank test between patients with a high vs. a low value of the CD8A/CSF1 ratio (binarization at the median). Survival analyses were performed using R 4.0.0 and the survival_3.1-12, survminer_0.4.7, and ggplot2 packages. For all the experiments, P values of 0.05, 0.01, 0.001 or 0.0001 were considered statistically significant, as designated by *, **, ***, ****, respectively.

## Acknowledgments

We would like to thank Dr. Amir Grau of the Technion-Israel Institute of Technology for assistance with the CyTOF experiments and Dr. Shira Ovadia for assistance in the animal facility.

## Author contributions

Conceptualization-ME, MP: Methodology-MP, JZ, SJ, AS, JB, LM, ON, MB, LC, KY, ST, BR, LB, AM, MS: Investigation-MP, JZ, SJ: Visualization-MP, ME: Funding acquisition-ME, MS: Project administration-ME, LC: Supervision-ME: Writing – original draft: ME, MP: Writing – review & editing-MP, SJ, LC, JF, IC, TC, IA, OD, BJ, DK, EV, MS, YC, JH, L.GT.M, PS, ME

## Data and materials availability

Raw RNA-seq data will be deposited at the NCBI gene expression Omnibus. All data are available in the main text or the supplementary materials. The raw data will be uploaded as a Source data. Any data and materials that can be shared will be released via a material transfer agreement.

---

**Supplementary Figure S1.**
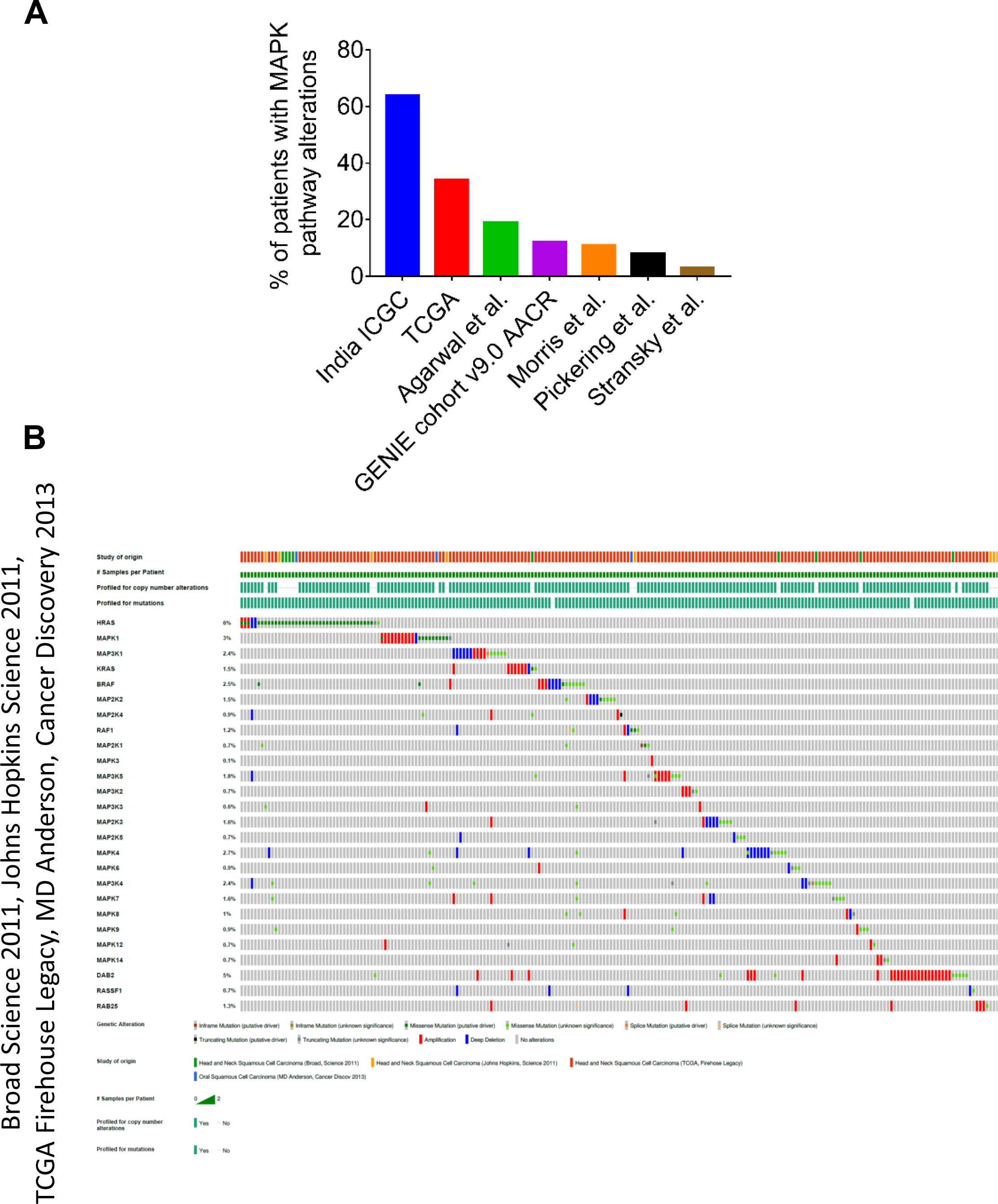
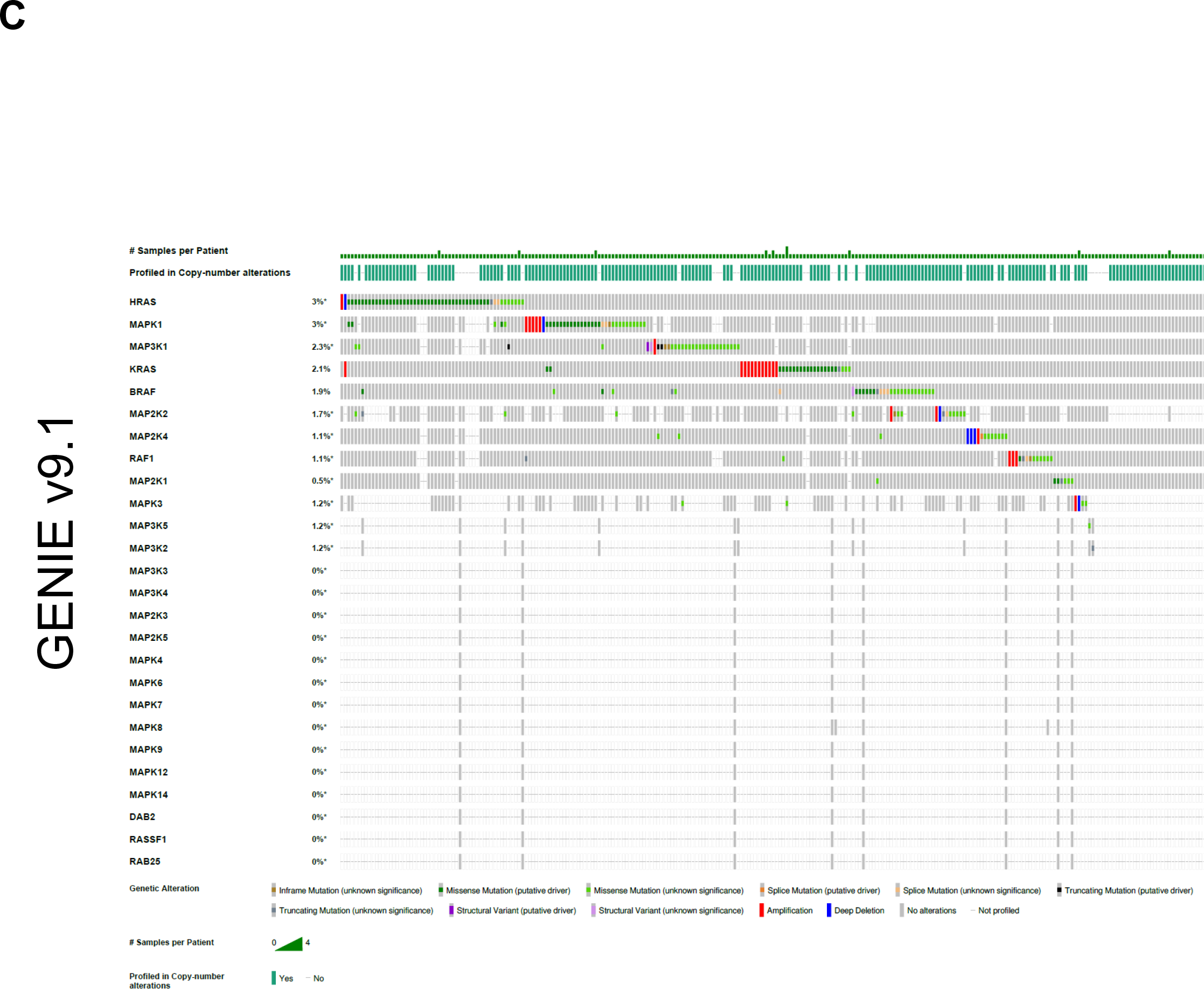
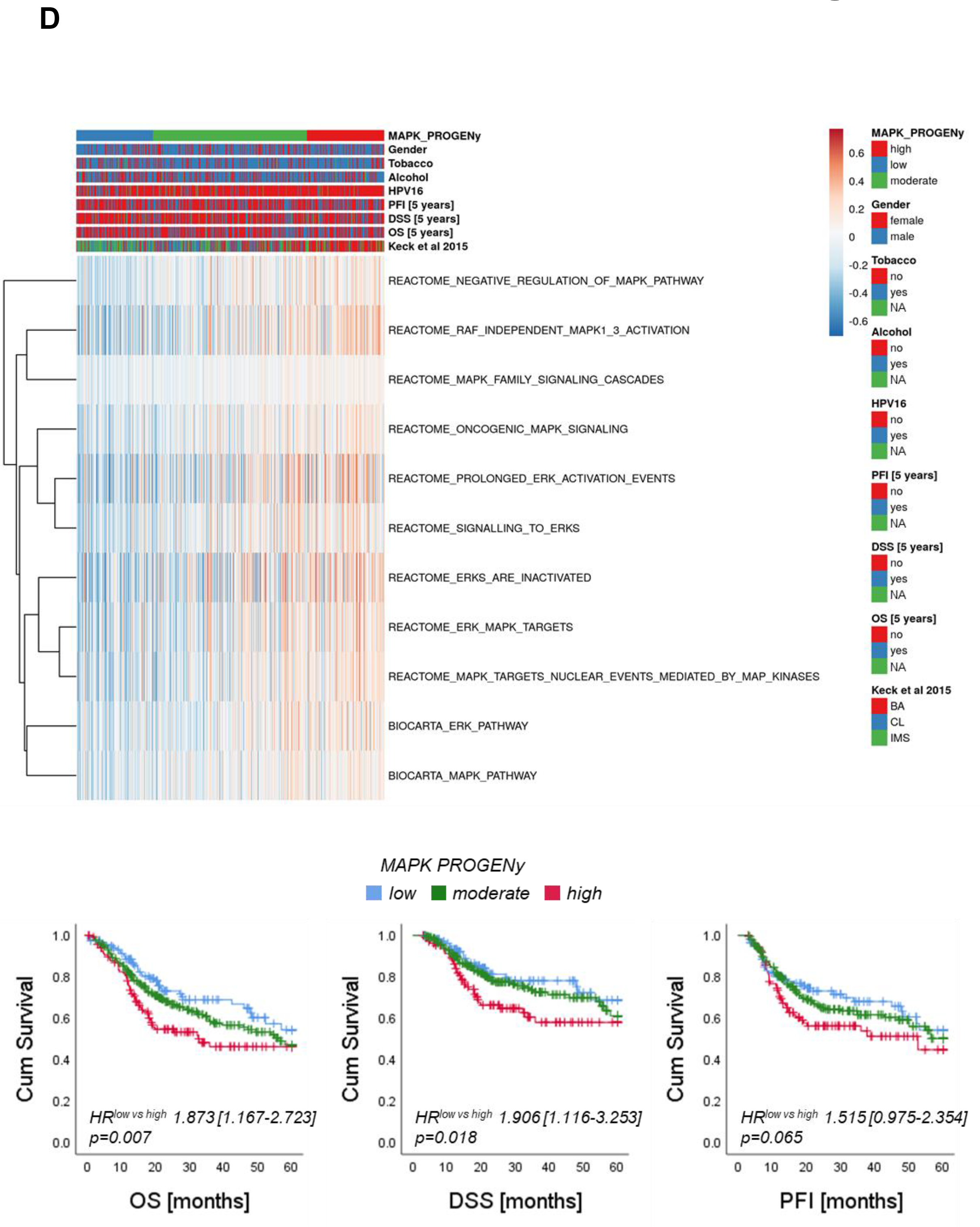
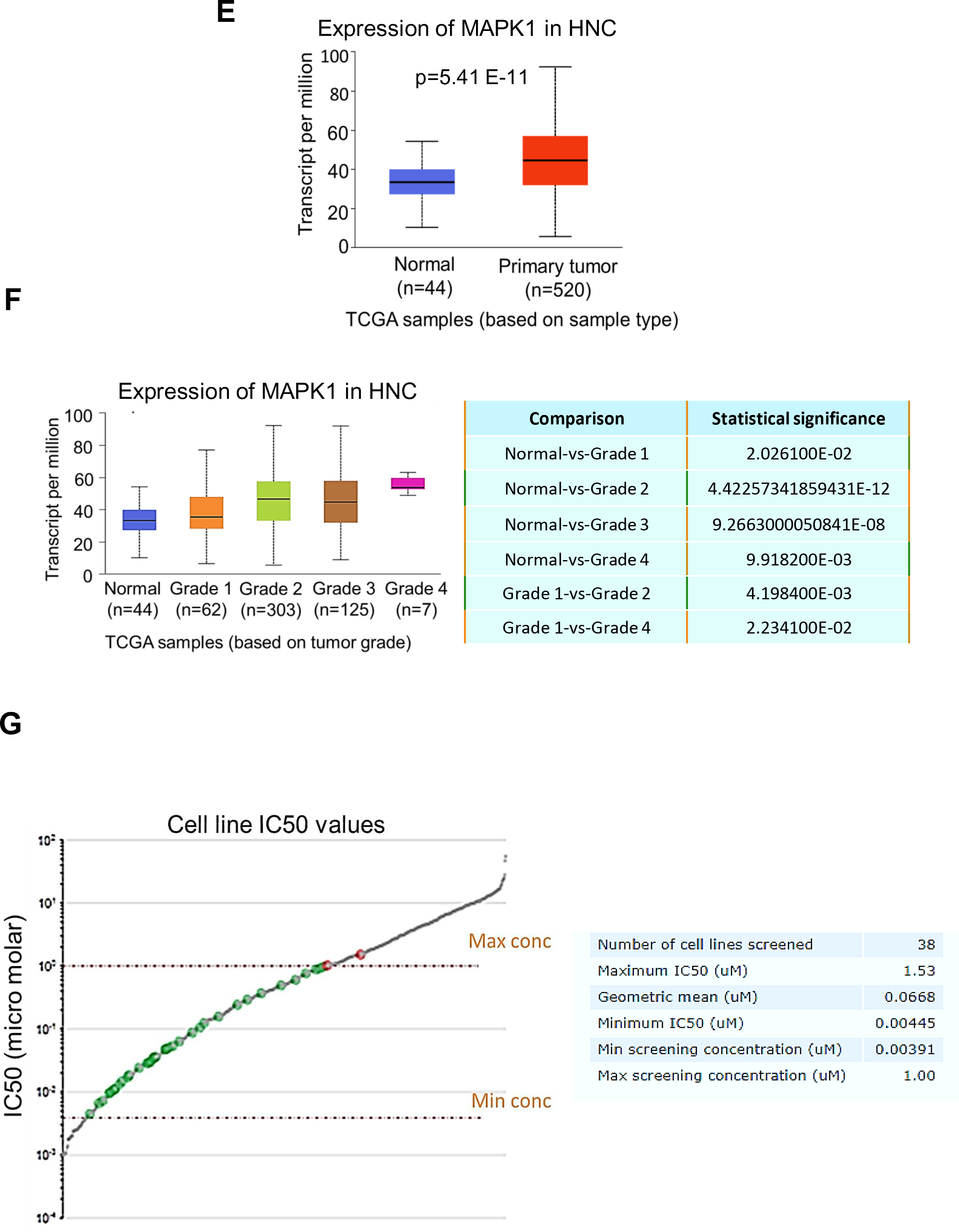
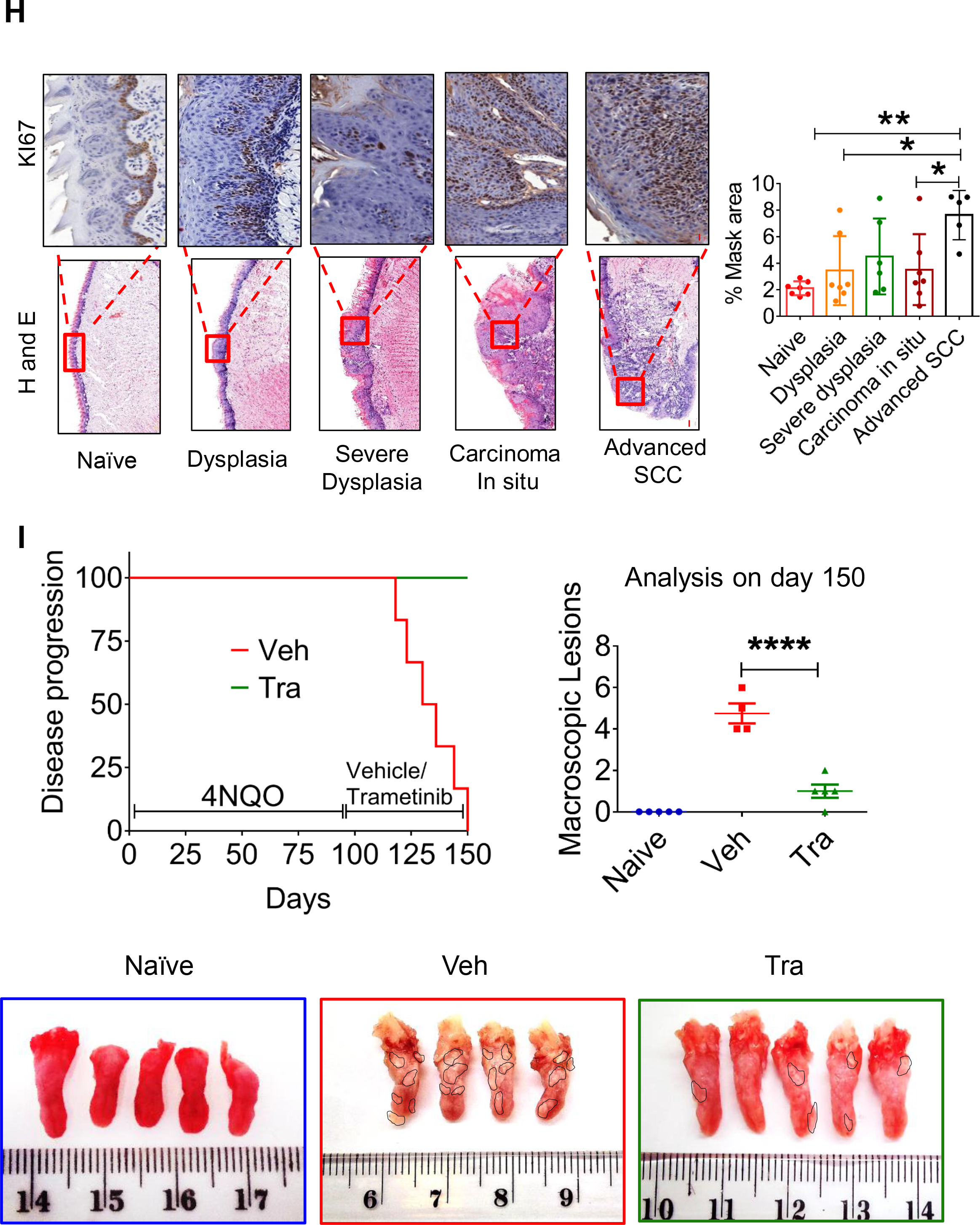
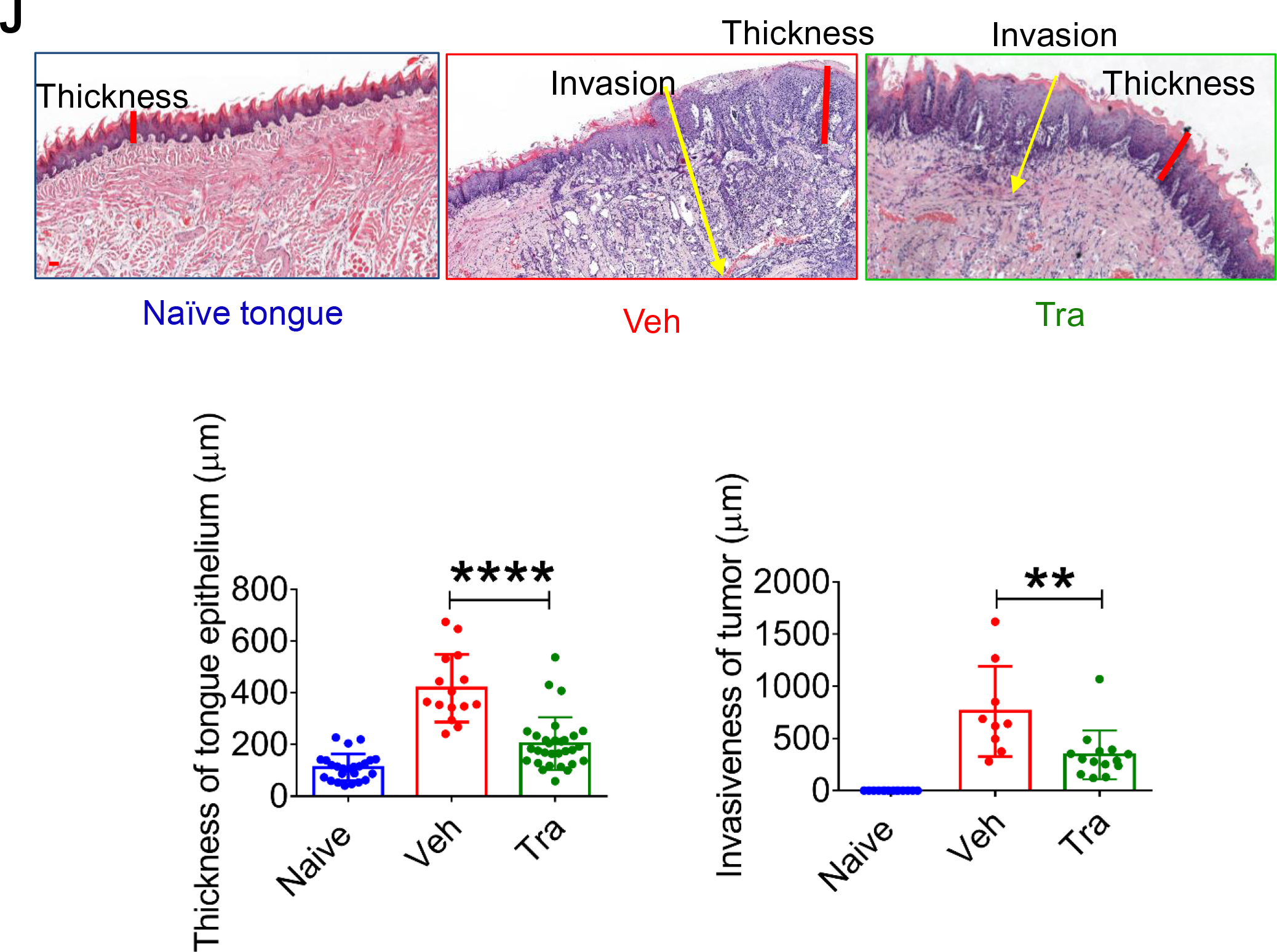
**(A)** Genomics analysis of multiple HNC cohorts showing the percentage of patients with alterations in genes of the MAPK pathway. (**B)** MAPK mutated genes in HNC patients – joint analysis of five cohorts of whole-exome sequencing. (**C)** MAPK mutated genes in HNC patients – analysis of the GENIE cohort. (**D)** Top – Heatmap illustrates GSVA score distribution for selected MAPK-related gene sets among tumors of TCGA-HNSC ranked according to MAPK activity inferred with the PROGENy algorithm. NA, not available. Bottom – Kaplan-Meier plots for five-year overall survival (OS; left), disease specific survival (DSS; middle) and progression-free intervals (PFI; right) for tumors with low, moderate and high MAPK pathway activity according to the PROGENy algorithm. (**E)** MAPK1 expression in normal tissue and in primary HNC tumors (TCGA database). (**F)** MAPK1 expression in different pathological grades of HNC (TCGA database); the table on the right presents the p values. (**G)** Analysis of the Cancer Cell Line Encyclopedia (CCLE) and Sanger cell line datasets showing the sensitivity of 38 cell lines to the MEK1/2 inhibitor, trametinib. (**H)** IHC images (top) of Ki67 staining and H&E images (bottom) showing various stages of oral (tongue) carcinogenesis induced by 4NQO in C57BL/6J mice. The percentage of mask area and statistics are also shown. Scale bars: 20 µm (top); 200 µm (bottom). (**I)** Top panel – disease progression of 4NQO-induced oral cancer (left) and quantification of macroscopic lesions (right). Bottom panel – Photographs showing the macroscopic lesions in the tongues of naïve, vehicle-exposed, and trametinib-treated mice after 150 days of the experiment. (**J)** H&E images and statistics for the tongues showing the thickness of the margins and invasion of the tumors (scale bars: 100 µm). For statistics, an unpaired 2-sided t-test or one-way ANOVA was performed. \**p* < 0.05; \*\**p* < 0.01; \*\*\**p* < 0.001, \*\*\*\**p* < 0.0001 were considered statistically significant. Tra - trametinib, Veh - vehicle.

**Supplementary Figure S2.**
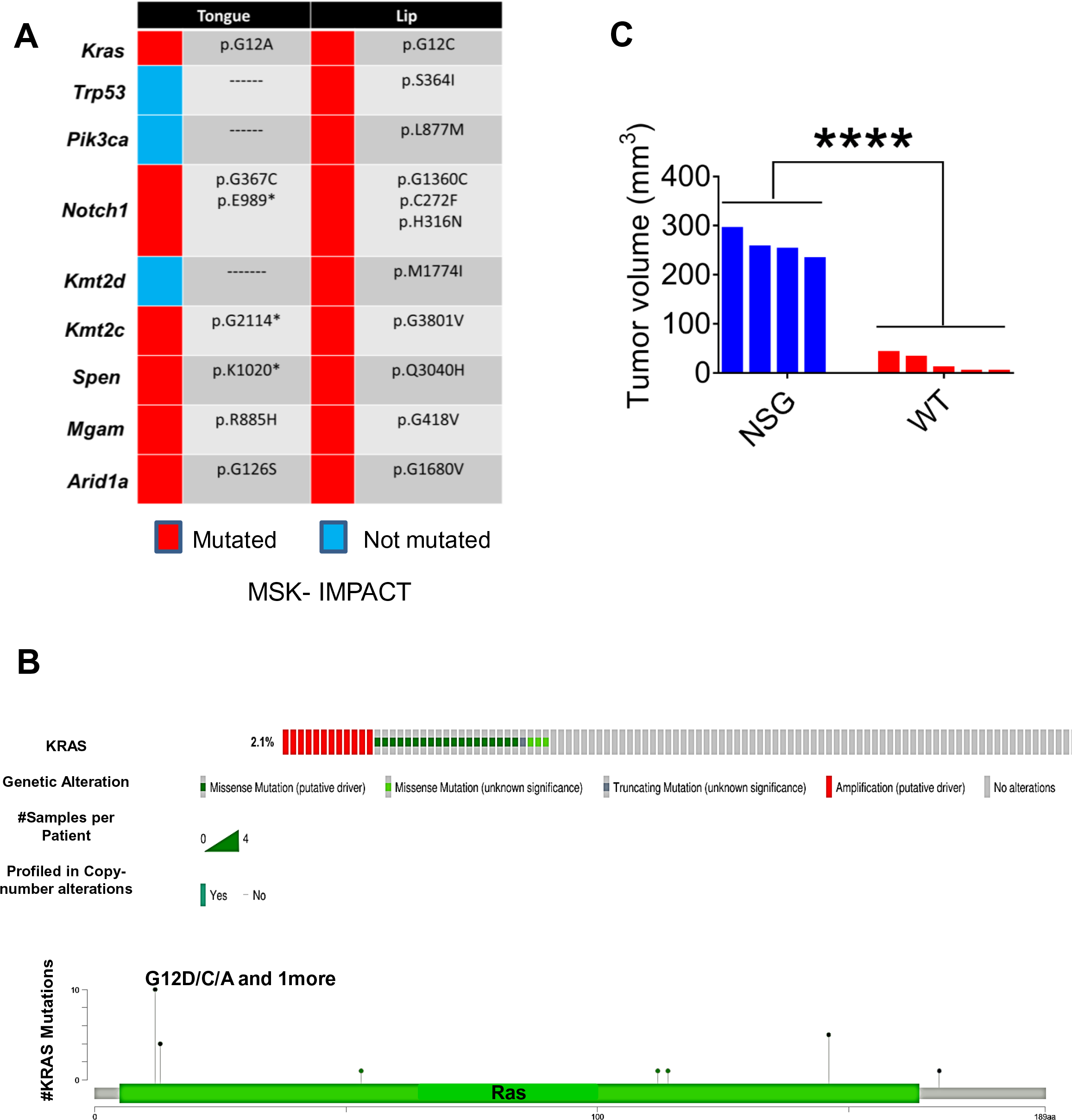
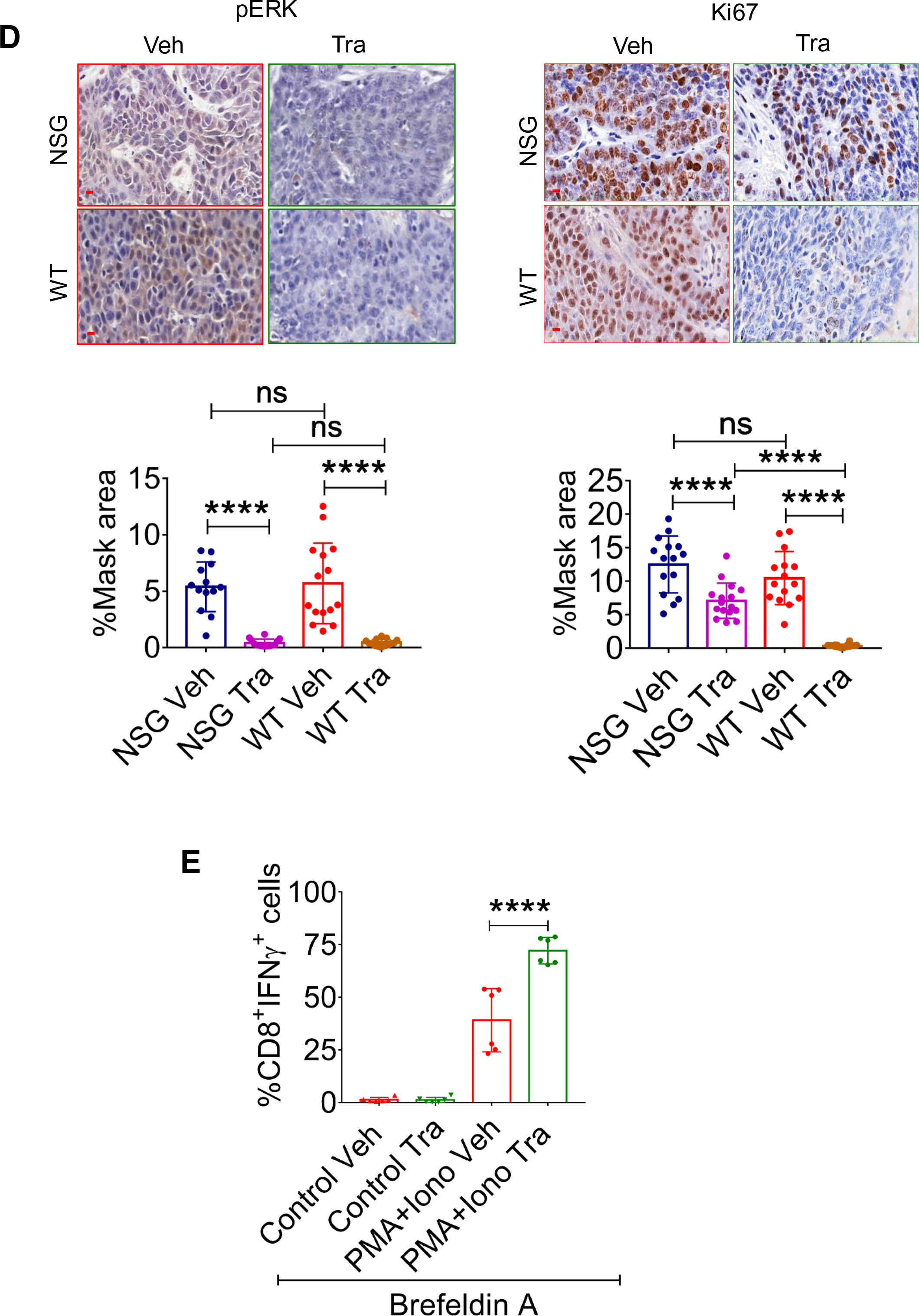
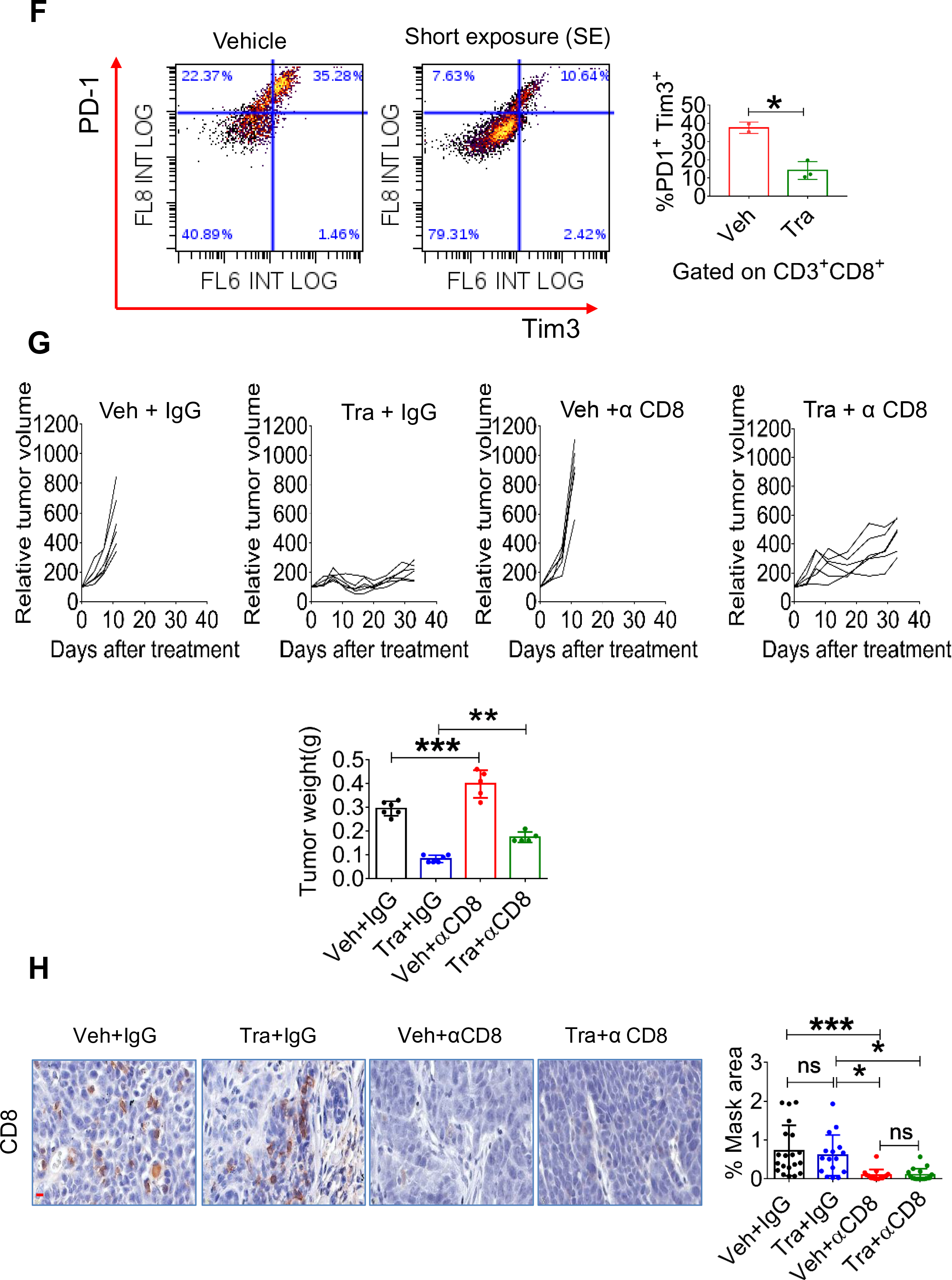
**(A)** Key mutations in 4NQO-T and 4NQO-L cell lines found in MSK-IMPACT genome sequencing. (**B)** Hot spot mutations of KRAS in HNC, extracted from cBioportal, GENIE cohort. (**C)** Tumor volume changes after 20 days of treatment with trametinib compared to vehicle in NSG and WT mice. (**D)** IHC staining of pERK (top left) and Ki67 (top right) in NSG and WT mice. Quantification is shown in the bottom panel (scale bars: 20 µm; inset 10 µm). (**E)** Intracellular staining of IFNγ in CD8^+^ T cells isolated from the vehicle-treated or trametinib SE-treated (5 days) mice. Graph showing the percentage of CD8^+^IFNγ^+^ with or without activation with phorbol 12-myristate 13-acetate (PMA) and ionomycin (Iono). Brefeldin A was used as protein transport inhibitor. Results of two independent experiments are shown. **(F)** PD-1 and TIM3 levels in CD8^+^ T cells in 4NQO-L tumors treated with vehicle or a short exposure with trametinib. **(G)** Top – Tumor growth curves for four groups of mice: vehicle + IgG, trametinib + IgG, vehicle + anti-CD8 (αCD8), and trametinib + αCD8. Bottom – Volume of 4NQO-L tumors in WT mice treated with vehicle or trametinib with and without depletion of CD8^+^ T cells. (**H)** IHC staining and quantification of CD8^+^ T cell depletion experiment (scale bar: 20 µm). For statistics, an unpaired 2-sided t-test or one-way ANOVA was performed. \**p* < 0.05; \*\**p* < 0.01; \*\*\**p* < 0.001, \*\*\*\**p* < 0.0001 were considered statistically significant. Tra - trametinib, Veh - vehicle.

**Supplementary Figure S3.**
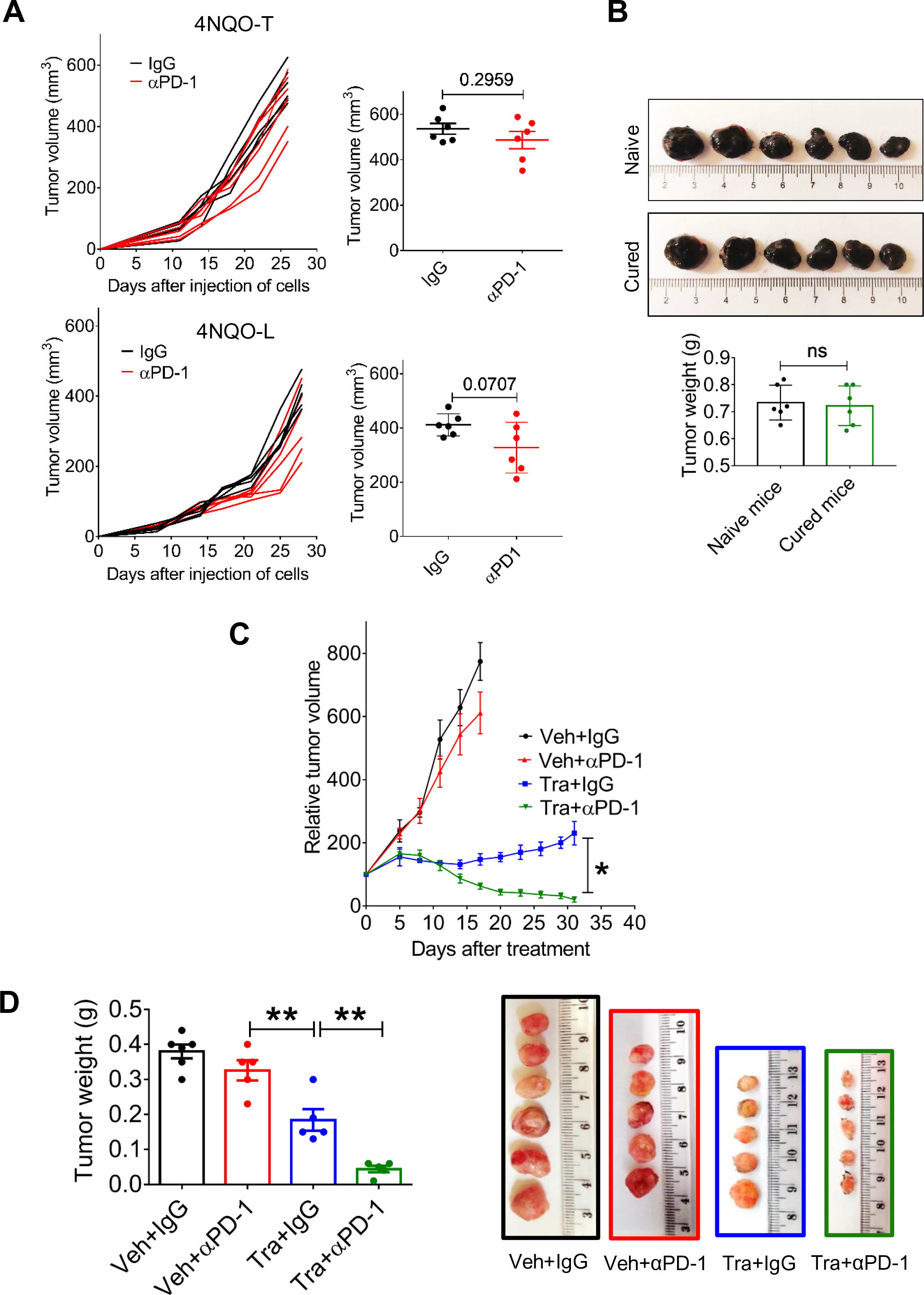
**(A)** Tumor growth curves and volumes at the experiment endpoint for 4NQO-L and 4NQO-T tumors in WT mice treated with αPD-1 or IgG. B16 tumors in naïve and cured mice and tumor weights in grams (bar diagram). Relative volumes of 4NQO-L tumors in WT mice treated as indicated. (**D)** Weights of 4NQO-L tumors and images of the tumors at the end of the experiment (day 31). For statistics, an unpaired 2-sided *t*-test or one-way ANOVA was performed. \**p* < 0.05, \*\**p* < 0.01, \*\*\**p* < 0.001, and \*\*\*\**p*< 0.0001 were considered statistically significant. ns - not significant. Tra - trametinib, Veh - vehicle.

**Supplementary Figure S4.**
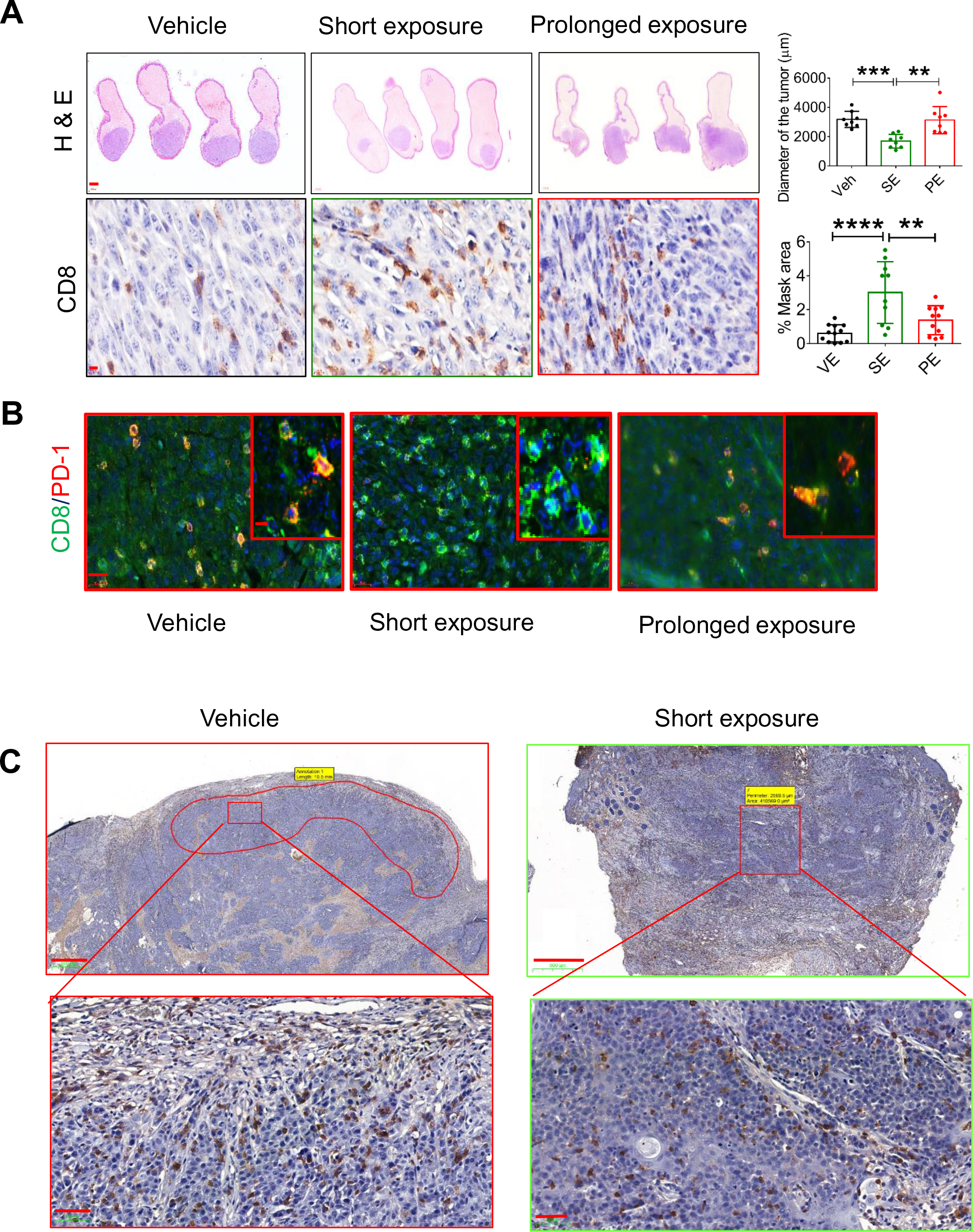
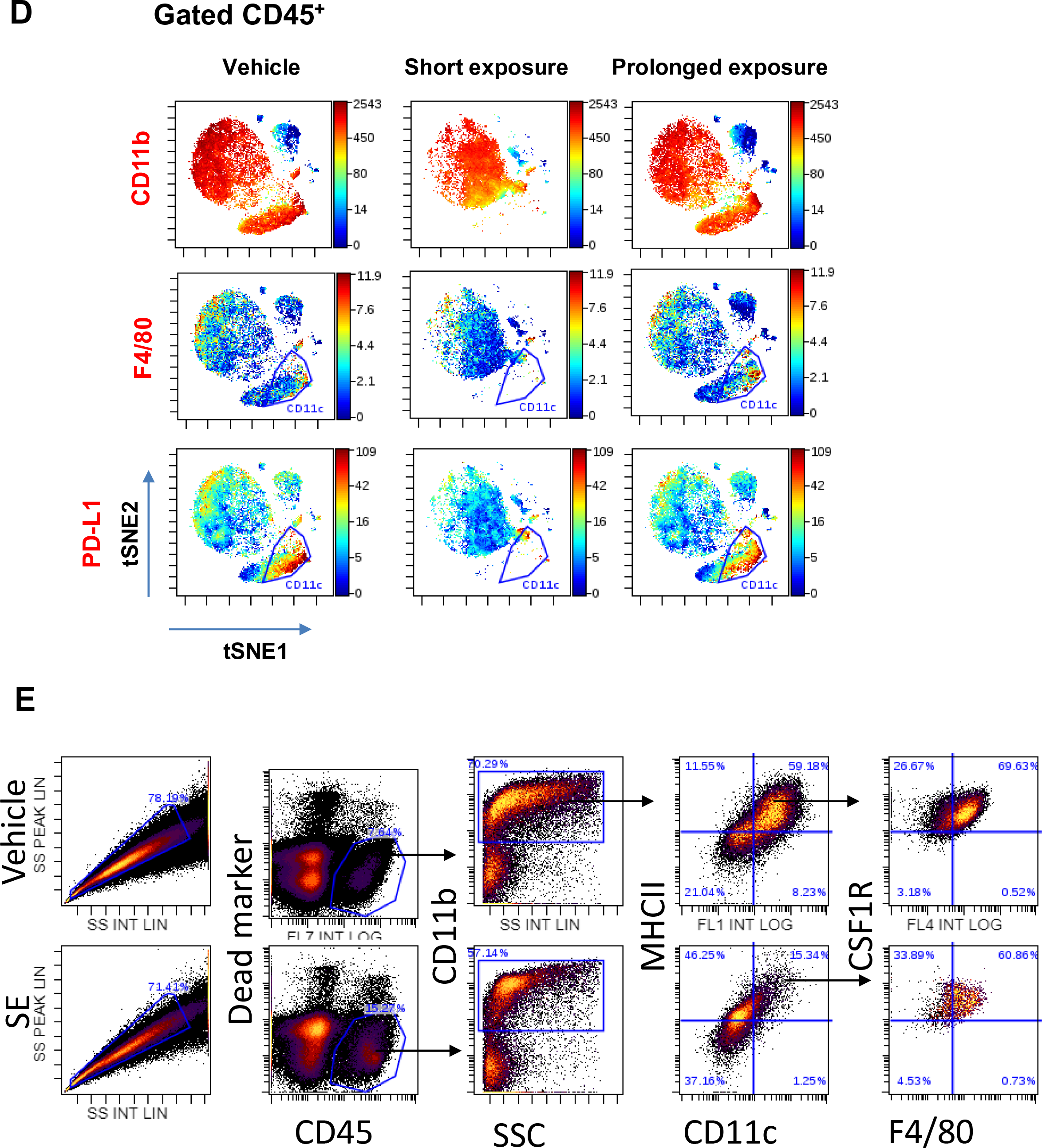
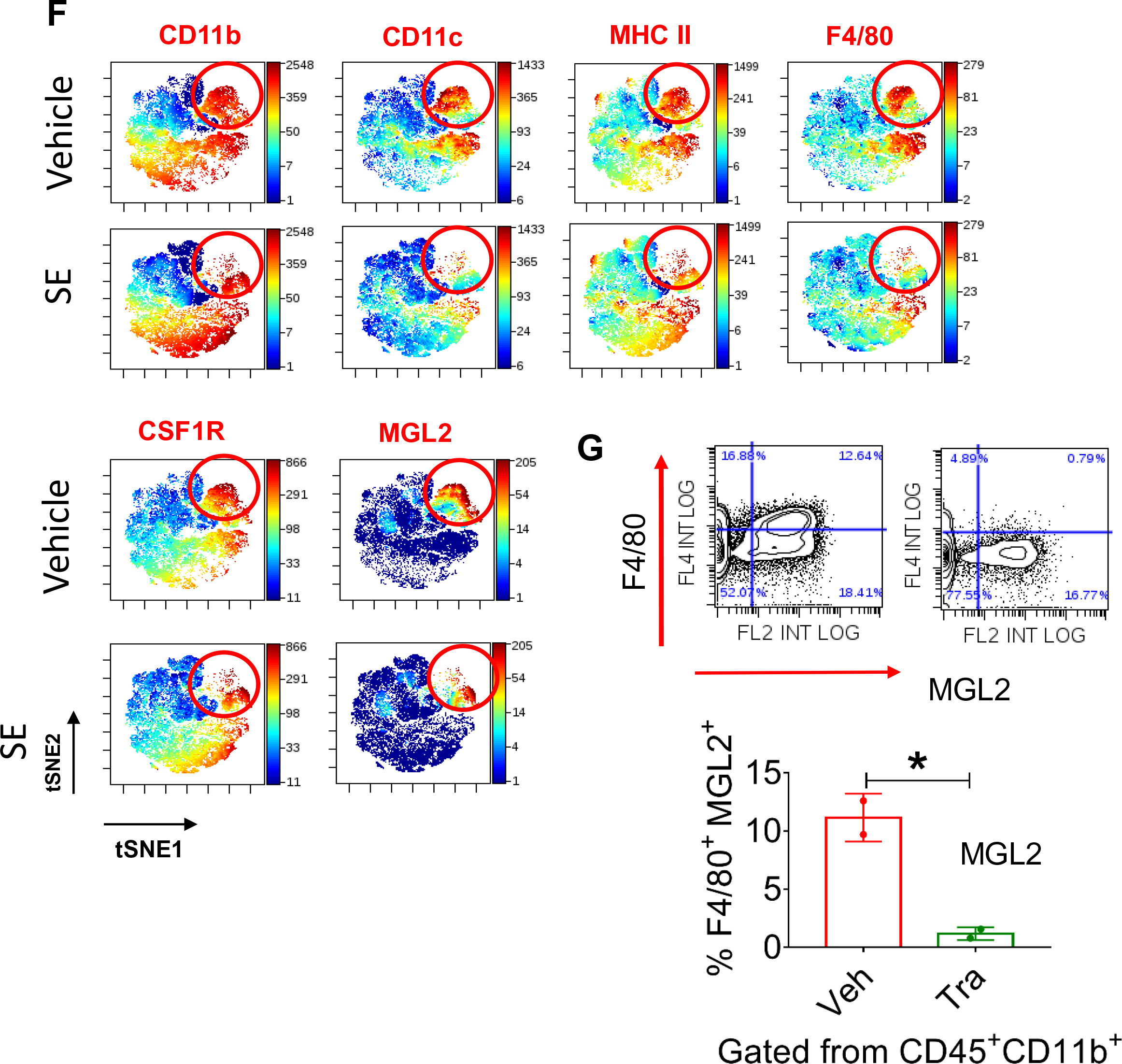
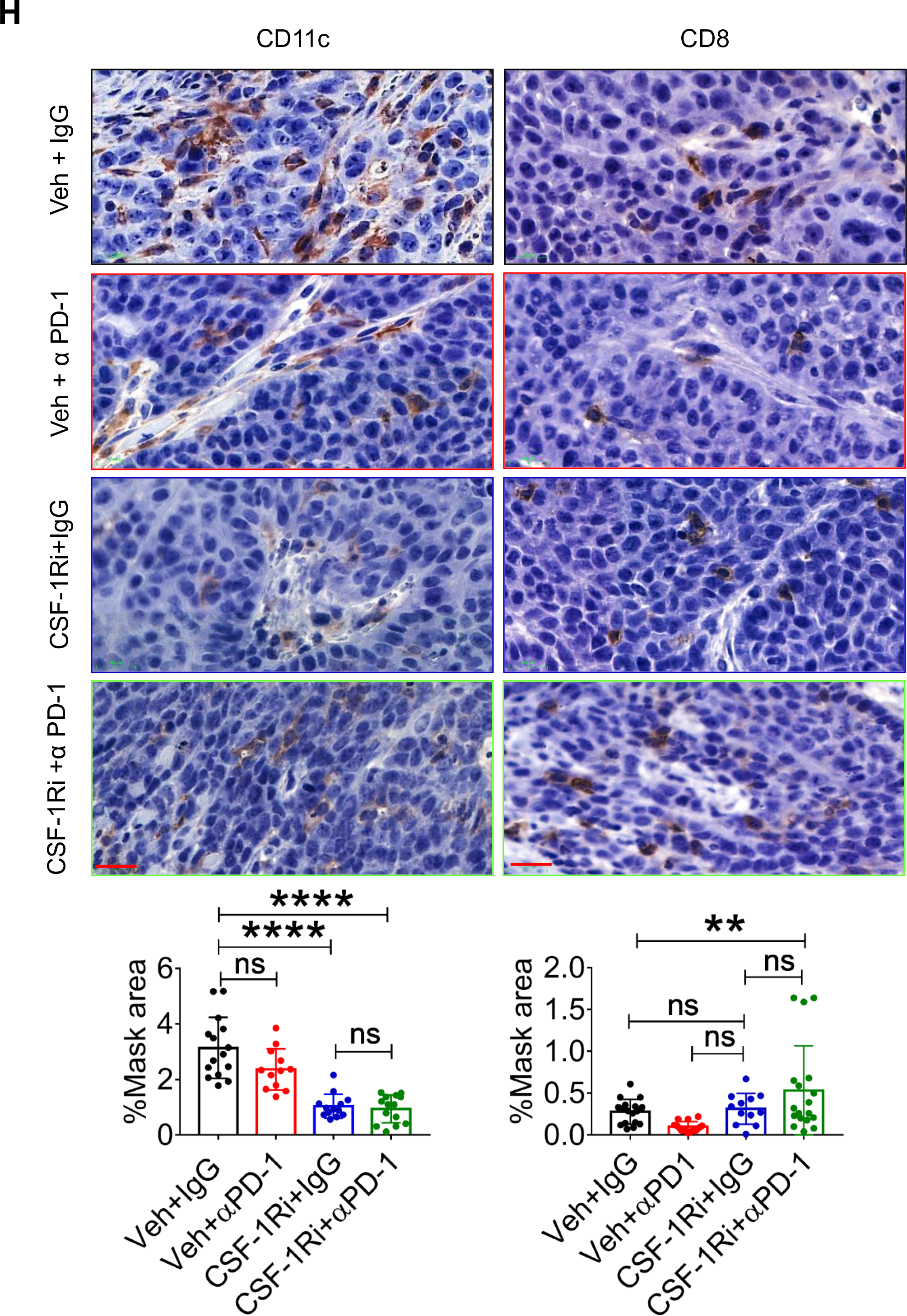
**(A)** Top – H&E stained images of tongues injected with 4NQO-T cells after exposure to vehicle, SE (7 days) trametinib or PE (25 days) trametinib (scale bars: 2000 µm). Bottom – IHC images and quantification of CD8^+^ T cells in 4NQO-T tumors after treatment with trametinib for 7 and 25 days (scale bars: 20 µm). (**B)** IF co-staining (OPAL) of CD8 (green) and PD1 (red) and merge (yellow) of 4NQO-T tumors treated as indicated (scale bars: 50 µm; inset 10 µm). (**C)** Pattern of infiltration of CD8^+^ T cells in the vehicle-treated and SE trametinib-treated lip tissue samples (Scale bars: 500 µm; enlargements 50 µm). (**D)** viSNE plots of the CyTOF data showing CD11b, F4/80 and PD-L1 expression in CD45^+^ cells from 4NQO-T tumors treated with vehicle, SE (5 days) trametinib, or PE (33 days) trametinib. (**E)** Flow cytometry analysis of CD45^+^ cells based on their CD11b, CD11c, MHCII CSF1R and F4/80 expression. **(F)** viSNE plots representing flow data of the myeloid populations CSF1R, CD11b, CD11c, MHCII, F4/80 and MGL2 (CD301b^+^) shown for 4NQO-L tumors treated with vehicle or trametinib for 5 days (SE). (**G)** Flow cytometry dot plot analysis of MGL2 (CD301b^+^) and F4/80 on CD45^+^CD11b^+^, when 4NQO-L tumors treated with a SE of trametinib or vehicle. (**H)** IHC analysis (top) and quantification (bottom) of CD11c and CD8 in tissues of 4NQO-L tumors treated with vehicle + IgG, vehicle + αPD-1, CSF-1Ri (CSF-1R inhibitor) + IgG, or CSF-1R + αPD-1 (scale bars: 20 µm). For statistics, an unpaired 2-sided *t*-test or one-way ANOVA was performed. \**p* < 0.05, \*\**p* < 0.01, \*\*\**p* < 0.001, and \*\*\*\**p* < 0.0001 were considered statistically significant. ns - not significant. Tra - trametinib, Veh - vehicle.

**Supplementary Figure S5.**
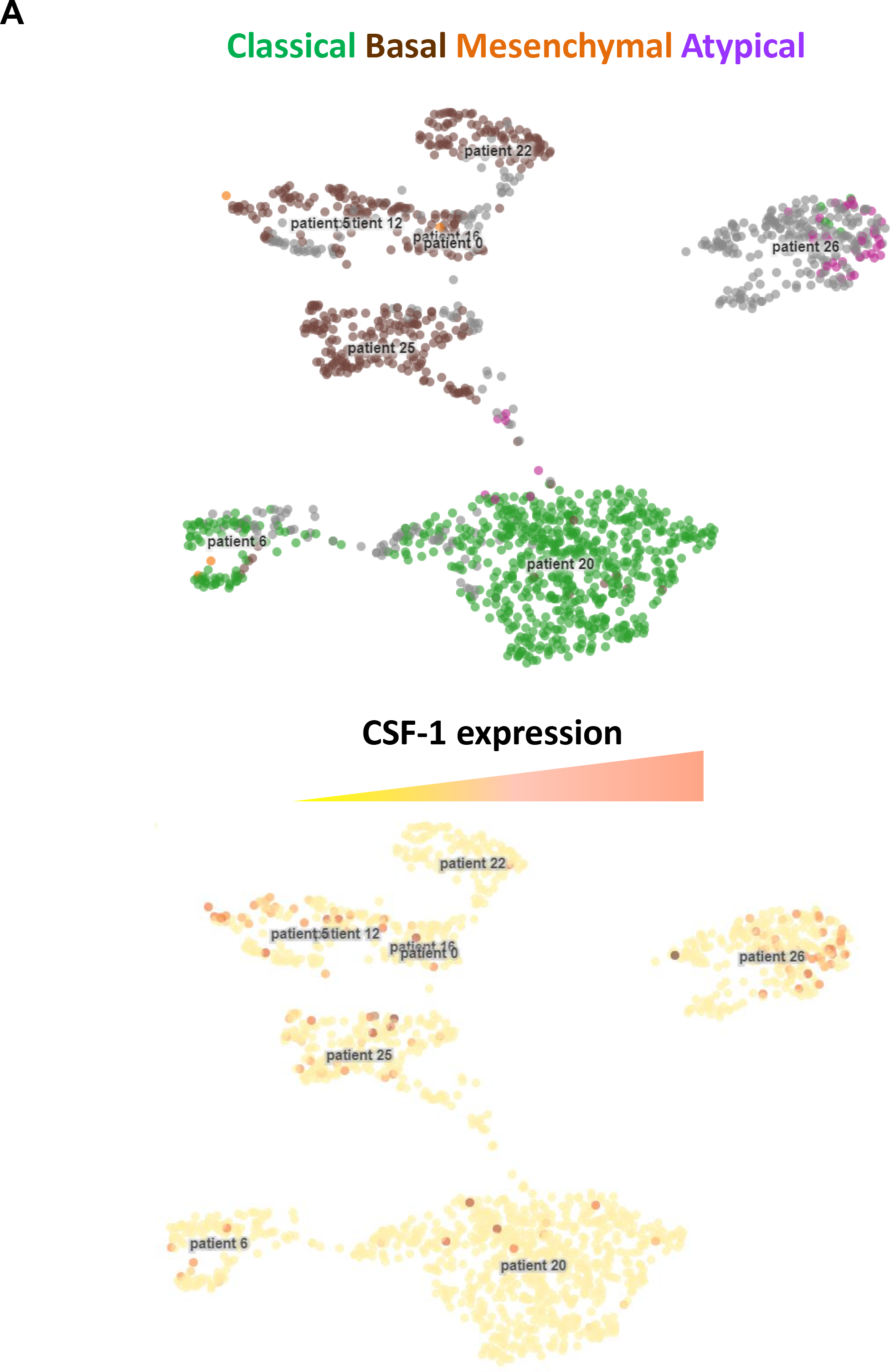
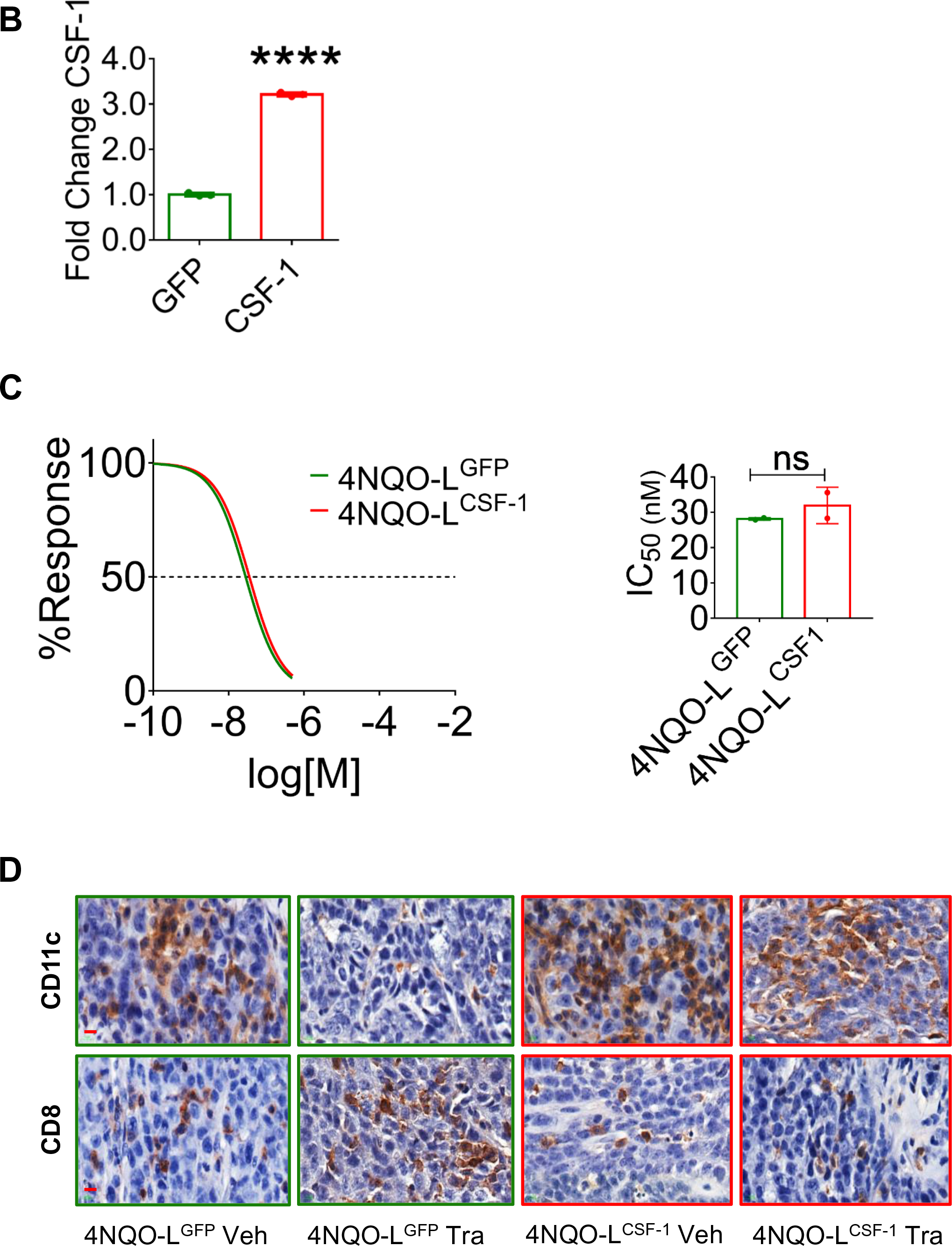
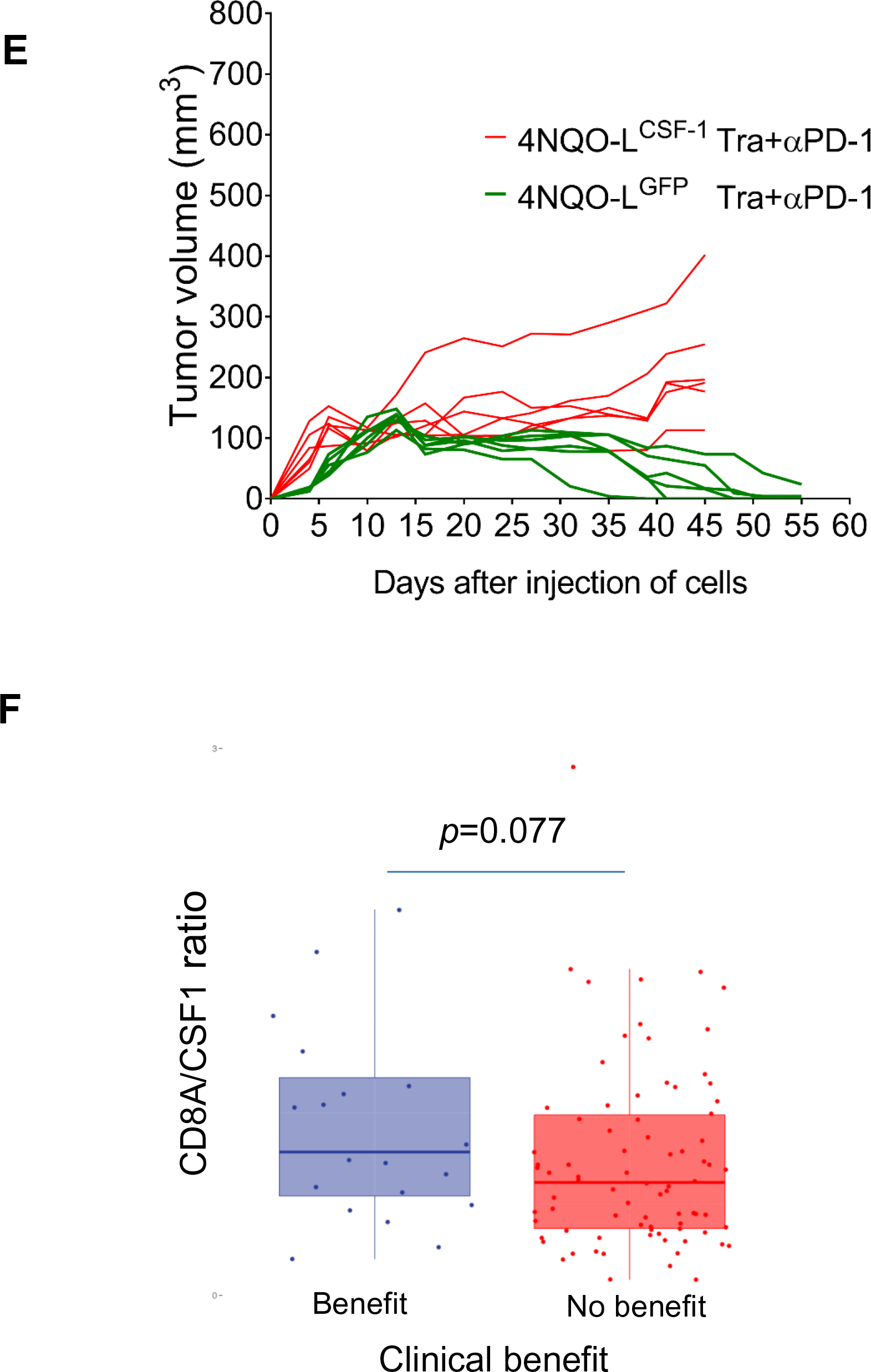
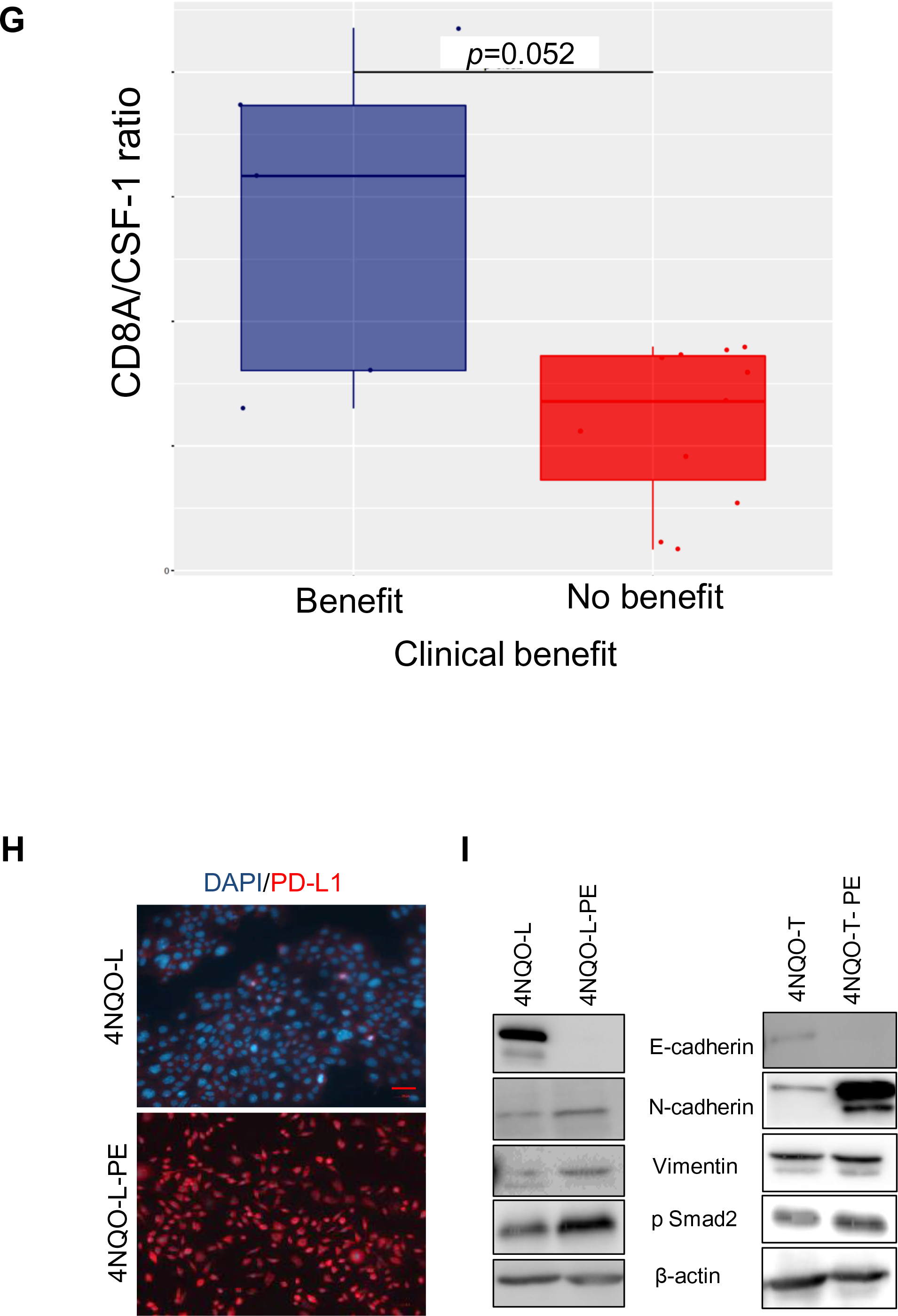
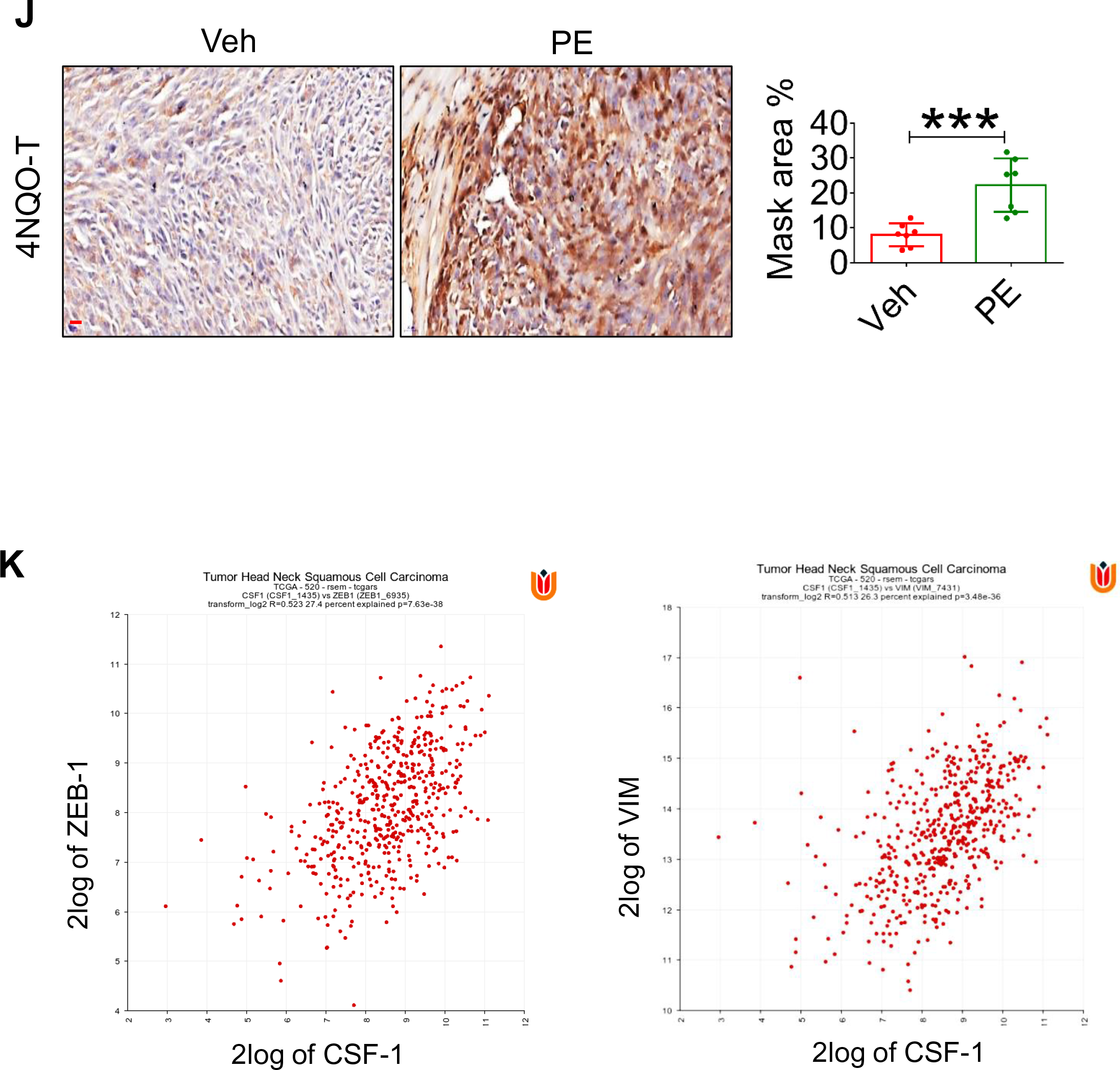
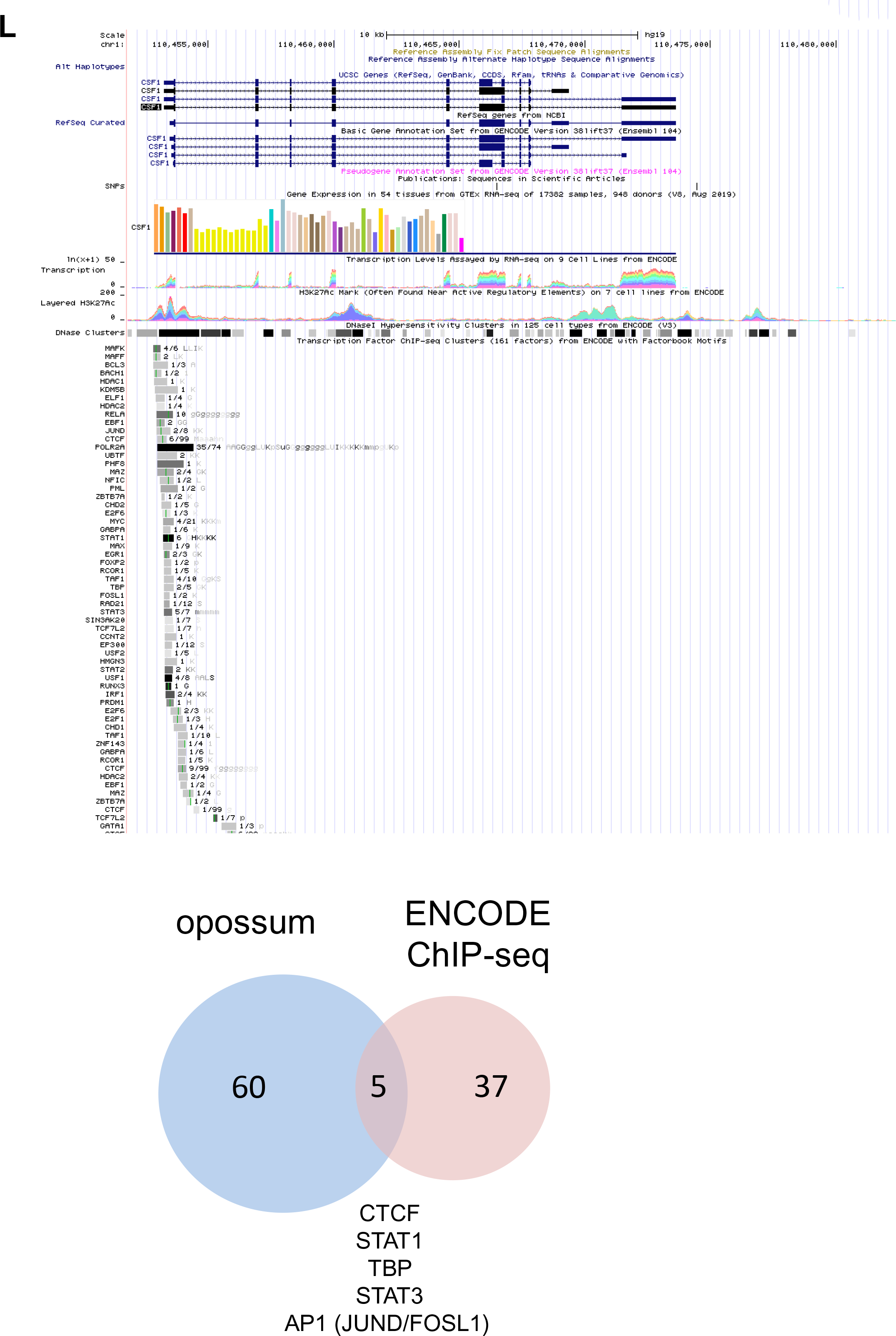
**(A)** Single-cell analysis of CSF-1 expression in tumor cells of HNC patients. (**B)** mRNA levels of CSF-1 in 4NQO-L-CSF-1 cells relative to 4NQO-L-GFP cells. **(C)** Dose-response curve (left) of trametinib in 4NQO-L-GFP and 4NQO-L-CSF-1 cells, and IC_50_ values obtained in two independent experiments (right). (**D)** IHC staining of CD8 and CD11c in 4NQO-L^GFP^ and 4NQO-L^CSF-1^ tumors treated with trametinib for 5 days (SE) (scale bars: 50 µm). (**E)** Growth curves of single mice with 4NQO-L^GFP^ or 4NQO-L^CSF-1^ tumors treated with a combination of trametinib and αPD-1 (n = 6). (**F)** Boxplots showing the relationship between the CD8A/CSF1 ratio and the clinical benefits of αPD1/PD-L1 treatment in 102 patients with head and neck squamous cell carcinoma (HNSCC) from the CLB-IHN cohort (based on RECIST criteria, clinical benefit was defined as complete response, partial response and stable disease of at least 6 months). A Mann-Whitney-Wilcoxon non-parametric test was used to compare the value of the CD8A/CSF1 ratio between the two groups. **(G)** Boxplots showing the value of the CD8A/CSF1 ratio in a lung cancer cohort. (**H)** IF images showing the expression of PD-L1 in 4NQO-L and 4NQO-L-PE cells. (**I)** Western blot analysis showing the expression levels of E-cadherin, N-cadherin, vimentin, pSMAD2 and β-actin in 4NQO-L, 4NQO-L-PE, 4NQO-T and 4NQO-T-PE cells. (**J)** IHC images and quantification of PD-L1 expression in 4NQO-T tumors treated with vehicle and trametinib for 30 days (PE) (scale bar: 100 µm). **(K)** Correlation of expression of CSF1 with ZEB1 and VIM, using the R2: Genomics Analysis and Visualization Platform. (**L)** ENCODE analysis of the promoter of CSF-1 and cross sectioning analysis of upregulated signature of TF in 4NQOs-PE cells and the TFs shown in ENCODE. For statistics, an unpaired 2-sided t-test or one-way ANOVA was performed. \**p* < 0.05; \*\**p* < 0.01; \*\*\**p* < 0.001, \*\*\*\**p* < 0.0001 were considered statistically significant. Tra - trametinib, Veh - vehicle.

## References

1. Rodriguez-Viciana P et al. Cancer targets in the Ras pathway. In: Cold Spring Harbor Symposia on Quantitative Biology. Cold Spring Harbor Laboratory Press; 2005:461–467

2. Sebolt-Leopold JS, Herrera R. Targeting the mitogen-activated protein kinase cascade to treat cancer. Nat. Rev. Cancer 2004;4(12):937–947.

3. Huang C et al. Proteogenomic insights into the biology and treatment of HPV-negative head and neck squamous cell carcinoma. Cancer Cell 2021;39(3):361–379.e16.

4. Ngan HL et al. MAPK pathway mutations in head and neck cancer affect immune microenvironments and ErbB3 signaling [Internet]. Life Sci. alliance 2020;3(6). doi:10.26508/lsa.201900545

5. Leemans CR, Braakhuis BJM, Brakenhoff RH. The molecular biology of head and neck cancer [Internet]. Nat. Rev. Cancer 2011;11(1):9–22.

6. Vermorken JB et al. Platinum-Based Chemotherapy plus Cetuximab in Head and Neck Cancer [Internet]. N. Engl. J. Med. 2008;359(11):1116–1127.

7. Uppaluri R et al. Biomarker and tumor responses of oral cavity squamous cell carcinoma to trametinib: A phase II neoadjuvant window-of-opportunity clinical trial. Clin. Cancer Res. 2017;23(9):2186–2194.

8. Verma V et al. MEK inhibition reprograms CD8+ T lymphocytes into memory stem cells with potent antitumor effects [Internet]. Nat. Immunol. [published online ahead of print: 2020]; doi:10.1038/s41590-020-00818-9

9. Kuske M et al. Immunomodulatory effects of BRAF and MEK inhibitors: Implications for Melanoma therapy. Pharmacol. Res. 2018;136:151–159.

10. Ebert PJR et al. MAP Kinase Inhibition Promotes T Cell and Anti-tumor Activity in Combination with PD-L1 Checkpoint Blockade. Immunity 2016;44(3):609–621.

11. Argiris A, Karamouzis M V., Raben D, Ferris RL. Head and neck cancer. Lancet 2008;371(9625):1695–1709.

12. Curry JM et al. Tumor Microenvironment in Head and Neck Squamous Cell Carcinoma [Internet]. Semin. Oncol. 2014;41(2):217–234.

13. Cillo AR et al. Immune landscape of viral- and carcinogen-driven head and neck cancer. Immunity 2020;52(1):183.

14. Dagogo-Jack I, Shaw AT. Tumour heterogeneity and resistance to cancer therapies [Internet]. Nat. Rev. Clin. Oncol. 2018;15(2):81–94.

15. Kohli K, Pillarisetty VG, Kim TS. Key chemokines direct migration of immune cells in solid tumors [Internet]. Cancer Gene Ther. 2021;1–12.

16. Burkholder B et al. Tumor-induced perturbations of cytokines and immune cell networks. Biochim. Biophys. Acta - Rev. Cancer 2014;1845(2):182–201.

17. Kang SH et al. Inhibition of MEK with trametinib enhances the efficacy of anti-PD-L1 inhibitor by regulating anti-tumor immunity in head and neck squamous cell carcinoma. Oncoimmunology 2019;8(1). doi:10.1080/2162402X.2018.1515057

18. Ferris RL et al. Nivolumab for Recurrent Squamous-Cell Carcinoma of the Head and Neck [Internet]. N. Engl. J. Med. 2016;375(19):1856–1867.

19. Chandrashekar DS et al. UALCAN: A Portal for Facilitating Tumor Subgroup Gene Expression and Survival Analyses [Internet]. Neoplasia (United States) 2017;19(8):649–658.

20. Lepikhova T et al. Drug-sensitivity screening and genomic characterization of 45 hpV-negative head and neck carcinoma cell lines for novel biomarkers of drug efficacy. Mol. Cancer Ther. 2018;17(9):2060–2071.

21. Kanojia D, Vaidya MM. 4-Nitroquinoline-1-oxide induced experimental oral carcinogenesis [Internet]. Oral Oncol. 2006;42(7):655–667.

22. Lydiatt WM et al. Head and neck cancers-major changes in the American Joint Committee on cancer eighth edition cancer staging manual. CA. Cancer J. Clin. 2017;67(2):122–137.

23. Kang S-H et al. Inhibition of MEK with trametinib enhances the efficacy of anti-PD-L1 inhibitor by regulating anti-tumor immunity in head and neck squamous cell carcinoma. Oncoimmunology 2018;00(00):1–11.

24. Lee JW et al. The Combination of MEK Inhibitor With Immunomodulatory Antibodies Targeting Programmed Death 1 and Programmed Death Ligand 1 Results in Prolonged Survival in Kras/p53-Driven Lung Cancer [Internet]. J. Thorac. Oncol. 2019;14(6):1046–1060.

25. Liu L et al. The BRAF and MEK Inhibitors Dabrafenib and Trametinib: Effects on Immune Function and in Combination with Immunomodulatory Antibodies Targeting PD-1, PD-L1, and CTLA-4. Clin. Cancer Res. 2015;21(7):1639–51.

26. Choi H et al. Pulsatile MEK Inhibition Improves Anti-tumor Immunity and T Cell Function in Murine Kras Mutant Lung Cancer [Internet]. CellReports 2019;27:806–819.e5.

27. Allegrezza MJ et al. Trametinib drives T-cell-dependent control of KRAS-mutated tumors by inhibiting pathological myelopoiesis [Internet]. Cancer Res. 2016;76(21):6253–6265.

28. Badarni M et al. Repression of AXL expression by AP-1/JNK blockage overcomes resistance to PI3Ka therapy [Internet]. JCI Insight 2019;4(8).https://insight.jci.org/articles/view/125341. cited April 10, 2019

29. Cheng DT et al. Memorial Sloan Kettering-Integrated Mutation Profiling of Actionable Cancer Targets (MSK-IMPACT): A Hybridization Capture-Based Next-Generation Sequencing Clinical Assay for Solid Tumor Molecular Oncology.. J. Mol. Diagn. 2015;17(3):251–64.

30. Chatila T, Silverman L, Miller R, Geha R. Mechanisms of T cell activation by the calcium ionophore ionomycin.. J. Immunol. 1989;143(4).

31. Crawford TQ, Jalbert E, Ndhlovu LC, Barbour JD. Concomitant evaluation of PMA+ionomycin-induced kinase phosphorylation and cytokine production in T cell subsets by flow cytometry. Cytom. Part A 2014;85(3):268–276.

32. Chen J et al. Single-cell transcriptomics reveal the intratumoral landscape of infiltrated T-cell subpopulations in oral squamous cell carcinoma. Mol. Oncol. [published online ahead of print: 2021]; doi:10.1002/1878-0261.12910

33. Ma J et al. PD1Hi CD8+ T cells correlate with exhausted signature and poor clinical outcome in hepatocellular carcinoma. J. Immunother. Cancer 2019;7(1):331.

34. Sanmamed MF et al. A Burned-Out CD8 + T-cell Subset Expands in the Tumor Microenvironment and Curbs Cancer Immunotherapy . Cancer Discov. [published online ahead of print: 2021]; doi:10.1158/2159-8290.cd-20-0962

35. Kenkel JA et al. An immunosuppressive dendritic cell subset accumulates at secondary sites and promotes metastasis in pancreatic cancer. Cancer Res. 2017;77(15):4158–4170.

36. Holmgaard RB et al. Timing of CSF-1/CSF-1R signaling blockade is critical to improving responses to CTLA-4 based immunotherapy [Internet]. Oncoimmunology 2016;5(7). doi:10.1080/2162402X.2016.1151595

37. Zhu Y et al. CSF1/CSF1R blockade reprograms tumor-infiltrating macrophages and improves response to T-cell checkpoint immunotherapy in pancreatic cancer models [Internet]. Cancer Res. 2014;74(18):5057–5069.

38. Puram S V. et al. Single-Cell Transcriptomic Analysis of Primary and Metastatic Tumor Ecosystems in Head and Neck Cancer [Internet]. Cell 2017;171(7):1611–1624.e24.

39. Cho JH et al. Osimertinib for patients with non-small-cell lung cancer harboring uncommon EGFR mutations: A multicenter, open-label, phase II trial (KCSG-Lu15-09). In: Journal of Clinical Oncology. American Society of Clinical Oncology; 2020:488–495

40. Kwon AT, Arenillas DJ, Hunt RW, Wasserman WW. oPOSSUM-3: Advanced Analysis of Regulatory Motif Over-Representation Across Genes or ChIP-Seq Datasets. G3 Genes|Genomes|Genetics 2012;2(9):987.

41. Homet Moreno B, Mok S, Comin-Anduix B, Hu-Lieskovan S, Ribas A. Combined treatment with dabrafenib and trametinib with immune-stimulating antibodies for BRAF mutant melanoma. Oncoimmunology 2016;5(7):e1052212.

42. Frederick DT et al. BRAF inhibition is associated with enhanced melanoma antigen expression and a more favorable tumor microenvironment in patients with metastatic melanoma. Clin. Cancer Res. 2013;19(5):1225–1231.

43. Ahn R, Ursini-Siegel J. Clinical Potential of Kinase Inhibitors in Combination with Immune Checkpoint Inhibitors for the Treatment of Solid Tumors. [Internet]. Int. J. Mol. Sci. 2021;22(5):1–23.

44. Kelly RJ. Dabrafenib and trametinib for the treatment of non-small cell lung cancer. Expert Rev. Anticancer Ther. 2018;18(11):1063–1068.

45. Biswas SK, Mantovani A. Macrophage plasticity and interaction with lymphocyte subsets: Cancer as a paradigm. Nat. Immunol. 2010;11(10):889–896.

46. Veglia F, Sanseviero E, Gabrilovich DI. Myeloid-derived suppressor cells in the era of increasing myeloid cell diversity [Internet]. Nat. Rev. Immunol. 2021; doi:10.1038/s41577-020-00490-y

47. Pixley FJ, Stanley ER. CSF-1 regulation of the wandering macrophage: Complexity in action [Internet]. Trends Cell Biol. 2004;14(11):628–638.

48. Neubert NJ et al. T cell-induced CSF1 promotes melanoma resistance to PD1 blockade [Internet]. Sci. Transl. Med. 2018;10(436). doi:10.1126/scitranslmed.aan3311

49. Pyonteck SM et al. CSF-1R inhibition alters macrophage polarization and blocks glioma progression [Internet]. Nat. Med. 2013;19(10):1264–1272.

50. Mok S et al. Inhibition of CSF-1 receptor improves the antitumor efficacy of adoptive cell transfer immunotherapy [Internet]. Cancer Res. 2014;74(1):153–161.

51. Beffinger M et al. CSF1R-dependent myeloid cells are required for NK-mediated control of metastasis [Internet]. JCI insight 2018;3(10). doi:10.1172/jci.insight.97792

52. Gyori D et al. Compensation between CSF1R+ macrophages and Foxp3+ Treg cells drives resistance to tumor immunotherapy [Internet]. JCI insight 2018;3(11). doi:10.1172/jci.insight.120631

53. Peranzoni E et al. Macrophages impede CD8 T cells from reaching tumor cells and limit the efficacy of anti–PD-1 treatment [Internet]. Proc. Natl. Acad. Sci. U. S. A. 2018;115(17):E4041–E4050.

54. Sun L et al. Inhibiting myeloid-derived suppressor cell trafficking enhances T cell immunotherapy [Internet]. JCI Insight 2019;4(7). doi:10.1172/jci.insight.126853

55. Jagadeeshan S et al. Adaptive Responses to Monotherapy in Head and Neck Cancer: Interventions for Rationale-Based Therapeutic Combinations. Trends in Cancer 2019;5(6). doi:10.1016/j.trecan.2019.04.004

56. Wagner S et al. Suppression of interferon gene expression overcomes resistance to MEK inhibition in KRAS-mutant colorectal cancer [Internet]. Oncogene 2019;38(10):1717–1733.

57. Mudianto T et al. Yap1 Mediates Trametinib Resistance in Head and Neck Squamous Cell Carcinomas[published online ahead of print: 2021]; doi:10.1158/1078-0432.CCR-19-4179

58. Segrelles C, Paramio JM, Lorz C. The transcriptional co-activator YAP: A new player in head and neck cancer [Internet]. Oral Oncol. 2018;86:25–32.

59. Kudo-Saito C, Shirako H, Takeuchi T, Kawakami Y. Cancer Metastasis Is Accelerated through Immunosuppression during Snail-Induced EMT of Cancer Cells [Internet]. Cancer Cell 2009;15(3):195–206.

60. Dongre A et al. Direct and Indirect Regulators of Epithelial-Mesenchymal Transition (EMT)-mediated Immunosuppression in Breast Carcinomas [Internet]. Cancer Discov. 2020;CD-20-0603.

61. Su S et al. A Positive feedback loop between mesenchymal-like cancer cells and macrophages is essential to breast cancer metastasis. Cancer Cell 2014;25(5):605–620.

62. Chae YK et al. Epithelial-mesenchymal transition (EMT) signature is inversely associated with T-cell infiltration in non-small cell lung cancer (NSCLC) [Internet]. Sci. Rep. 2018;8(1). doi:10.1038/s41598-018-21061-1

63. da Silva SD et al. Co-overexpression of twist1-csf1 is a common event in metastatic oral cancer and drives biologically aggressive phenotype [Internet]. Cancers (Basel). 2021;13(1):1–19.

64. Peng DH et al. Collagen promotes anti-PD-1/PD-L1 resistance in cancer through LAIR1-dependent CD8+ T cell exhaustion [Internet]. Nat. Commun. 2020;11(1). doi:10.1038/s41467-020-18298-8

65. Wang G et al. The pan-cancer landscape of crosstalk between epithelial-mesenchymal transition and immune evasion relevant to prognosis and immunotherapy response. *npj Precis*. Oncol. 2021;5(1). doi:10.1038/s41698-021-00200-4

66. Haas L et al. Acquired resistance to anti-MAPK targeted therapy confers an immune-evasive tumor microenvironment and cross-resistance to immunotherapy in melanoma. *Nat*. Cancer 2021 2021;1– 16.

67. Cancer Genome Atlas NetworkParticipants are arranged by area of contribution T, by institution then. ARTICLE Comprehensive genomic characterization of head and neck squamous cell carcinomas. Nature 2014;517. doi:10.1038/nature14129

68. Kansy BA et al. PD-1 status in CD8+ T cells associates with survival and anti-PD-1 therapeutic outcomes in head and neck cancer [Internet]. Cancer Res. 2017;77(22):6353–6364.

69. K K, et al. Establishment of a Plasticity-Associated Risk Model Based on a SOX2- and SOX9-Related Gene Set in Head and Neck Squamous Cell Carcinoma. Mol. Cancer Res. 2021;molcanres.MCR-21-0066-A.2021.

70. Stransky N et al. The Mutational Landscape of Head and Neck Squamous Cell Carcinoma [Internet]. Science 2011;333(6046):1157.

71. Morris LGT et al. The molecular landscape of recurrent and metastatic head and neck cancers insights from a precision oncology sequencing platform [Internet]. JAMA Oncol. 2017;3(2):244–255.

72. Sweeney SM et al. AACR project genie: Powering precision medicine through an international consortium [Internet]. Cancer Discov. 2017;7(8):818–831.

73. Agrawal N et al. Exome sequencing of head and neck squamous cell carcinoma reveals inactivating mutations in NOTCH1. [Internet]. Science 2011;333(6046):1154–7.

74. Metsalu T, Vilo J. ClustVis: a web tool for visualizing clustering of multivariate data using Principal Component Analysis and heatmap. Nucleic Acids Res. 2015;43(Web Server issue):W566.

75. Schubert M et al. Perturbation-response genes reveal signaling footprints in cancer gene expression. Nat. Commun. 2018;9(1). doi:10.1038/S41467-017-02391-6

76. A M, et al. EGFR and PI3K Pathway Activities Might Guide Drug Repurposing in HPV-Negative Head and Neck Cancers.. Front. Oncol. 2021;11:678966–678966.

77. Liberzon A et al. The Molecular Signatures Database Hallmark Gene Set Collection. Cell Syst. 2015;1(6):417–425.

78. Subramanian A et al. From the Cover: Gene set enrichment analysis: A knowledge-based approach for interpreting genome-wide expression profiles. Proc. Natl. Acad. Sci. U. S. A. 2005;102(43):15545.

79. Yegodayev KM et al. TGF-Beta-Activated Cancer-Associated Fibroblasts Limit Cetuximab Efficacy in Preclinical Models of Head and Neck Cancer [Internet]. Cancers (Basel). 2020;12(2):339.

80. Remer E et al. CDK 4/6 Inhibition Overcomes Acquired and Inherent Resistance to PI3Kα Inhibition in Pre-Clinical Models of Head and Neck Squamous Cell Carcinoma. J. Clin. Med. 2020;9(10):3214.

81. Finck R et al. Normalization of mass cytometry data with bead standards. Cytom. Part A 2013;83 A(5):483–494.

82. Kotecha N, Krutzik PO, Irish JM. Web-based analysis and publication of flow cytometry experiments. Curr. Protoc. Cytom. 2010;Chapter 10(SUPPL. 53). doi:10.1002/0471142956.cy1017s53

83. Karabajakian A et al. Hyperprogression and impact of tumor growth kinetics after PD1/PDL1 inhibition in head and neck squamous cell carcinoma. Oncotarget 2020;11(18):1618–1628.

